# Early-adult methionine restriction reduces methionine sulfoxide and extends lifespan in *Drosophila*

**DOI:** 10.1101/2023.03.15.532514

**Authors:** Hina Kosakamoto, Fumiaki Obata, Junpei Kuraishi, Hide Aikawa, Rina Okada, Joshua N. Johnstone, Taro Onuma, Matthew D. W. Piper, Masayuki Miura

**Author notes:** Co-corresponding authors (F.O.), (M.M.). Co-first authors (Hina Kosakamoto, Fumiaki Obata).

## Abstract

Methionine restriction (MetR) extends lifespan in various organisms, but its mechanistic understanding remains incomplete. Whether MetR during a specific period of adulthood increases lifespan is not shown. In *Drosophila*, MetR is reported to extend lifespan only when amino acid levels are low. Here, by using an exome-matched holidic medium, we show that decreasing Met levels to 10% extends *Drosophila* lifespan with or without decreasing total amino acid levels. MetR during the first four weeks of adult life robustly extends lifespan. MetR induces the expression of *Methionine sulfoxide reductase A* (*MsrA*) in young flies, which reduces the oxidatively-damaged Met. *MsrA* induction is *foxo*-dependent and persists for two weeks after cessation of the MetR diet. Loss of *MsrA* attenuates lifespan extension by early-adult MetR. Our study highlights the age-dependency of the organismal response to specific nutrient and suggests that nutrient restriction at a particular period of life is sufficient for healthspan extension.

## Introduction

Lifelong dietary restriction (DR) is a robust intervention for extending lifespan in various model organisms^1–3^. Many epidemiological and biological data suggest that DR can also be beneficial in humans^4,5^. However, this may depend on age. Lower dietary protein intake can reduce overall mortality in those 65 years or younger, but not in those older than 65 years^6^. A large-scale lifespan analysis using eight hundred mice demonstrated that DR only in the period of 3-24 months can extend lifespan, although the mortality rate is acutely increased upon swapping diet restricted animals to normal chow^7^. In contrast, DR started at 24 months exerts a blunted transcriptome response, especially in the fat tissues, and results in a minor increase in lifespan, suggesting nutritional memory^7^.

*Drosophila melanogaster,* due to its accessibility to genetic and dietary manipulation, is a useful tool for ageing research^8,9^. In *Drosophila*, the effect of DR was reported to be reversible, hence swapping the DR to the fully-fed diet on day 18 almost completely reversed the decrease in mortality^10^. However, a high-sugar diet for the first two-to-three weeks in adulthood showed a long-lasting effect on gene expression and lifespan, suggesting the memory of the early adult diet also in *Drosophila*^11^. Like DR, intermittent fasting (IF) is another dietary regimen for extending longevity. A two-days fed: five-days fasted IF condition only in the first month is sufficient for extending *Drosophila* lifespan^12^. Brief rapamycin treatment in the first two weeks can increase female lifespan via prolonged intestinal autophagy^13^. These reports together raise the question of to what extent and by which mechanisms the organism is affected by the early-adult dietary condition.

Methionine (Met), an essential amino acid, is a dietary requirement for animals. Specific restriction of dietary Met has been reported to extend lifespan in rodents^14,15^. Subsequent studies demonstrated that Met restriction (MetR) can extend lifespan in many model organisms, including *Drosophila*^16–19^. It has been suggested in *Drosophila* that active Met metabolism is a key mechanism for longevity^20–23^. Although several possible mechanisms by which dietary Met regulates the healthspan of organisms have been implicated, how MetR remodels organismal physiology is not fully understood^24,25^. In addition, whether short-term MetR, especially in early adulthood, can influence the organismal lifespan has not been fully studied. In this study, we asked whether transient MetR can impact organismal lifespan in *Drosophila*. Using tissue-specific and single-cell RNAseq, we describe how early MetR alters transcriptome and tissue homeostasis.

## Results

### Methionine restriction robustly extends fly lifespan

In *Drosophila*, Met restriction (MetR) extends female lifespan, especially when total amino acid (AA) levels are reduced to 40% of the control^19^. First, we tested whether MetR in our laboratory condition could reproducibly extend lifespan. In our study, we utilised an exome-matched version of the holidic medium developed by Piper et al.^26^ and decreased all AA concentrations to 40% of the original recipe. For Met restriction (MetR), the Met concentration was further decreased to 10% of the control, which resulted in 0.16 mM Met in the MetR diet and 1.6 mM Met in the control diet. This resulted in a negligible (less than 1%) change in the total energy content of the diet from 95.36 kcal/L for the control, to 94.49 kcal/L for MetR. Under this condition, we observed an increase in the median lifespan of a wild-type Canton-S female of up to 34.5% (Fig. 1a). The effect of the MetR diet was also observed in the outbred strain *w^Dah^* (Fig. 1b), showing the robustness of the phenotype.

**Fig. 1.**
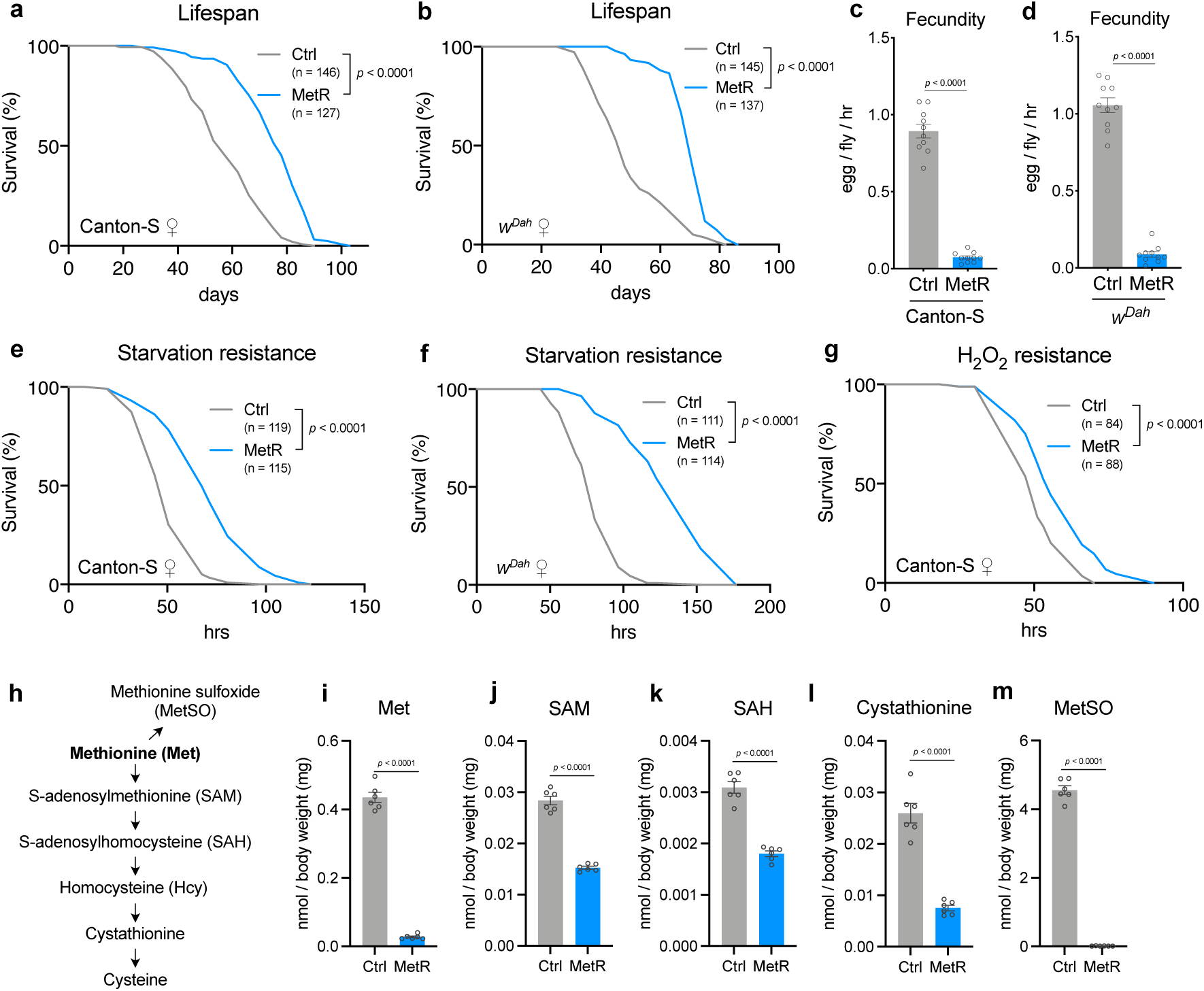
Methionine restriction decreases its metabolites and extends female lifespan. **a, b,** Lifespans of female Canton-S (**a**) and *w^Dah^* (**b**) flies fed with or without a methionine-restricted diet. Sample sizes (n) are shown in the figure. For the statistics, a log-rank test was used. **c, d,** Fecundity of Canton-S (**c**) and *w^Dah^*(**d**) flies fed with or without a methionine-restricted diet. n = 10. For the statistics, a two-tailed Student’s *t* test was used. **e, f,** Survivability of female Canton-S (**e**) or *w^Dah^*(**f**) flies upon complete starvation after feeding with or without a methionine-restricted diet for one week. Sample sizes (n) are shown in the figure. **g,** H_2_O_2_ resistance of female Canton-S flies after feeding with or without a methionine-restricted diet for one week. Sample sizes (n) are shown in the figure. **h,** Methionine metabolic pathway and its oxidation to methionine sulfoxide. **i-m,** Quantification of methionine metabolites and the oxidative product upon methionine restriction. n = 6. For the statistics, a two-tailed Student’s *t* test was used. For all graphs, the mean and SEM are shown. Data points indicate biological replicates.

Next, we quantified the fecundity (egg-laying) and the resistance to stressors. Flies fed with the MetR diet for one week showed a decrease in the number of eggs and a marked increase in starvation resistance in both Canton-S and *w^Dah^* (Fig. 1c-f). This phenotype is reminiscent of what has been observed in flies under dietary restriction (DR)^27^. It has been reported that DR increases starvation resistance via increased lipids in the gut^28^. As expected, we observed accumulation of lipid staining in the gut during MetR (Supplementary Fig. 1a, b). MetR also enhanced resistance to an oxidant, hydrogen peroxide, at least in Canton-S (Fig. 1g). Increased stress resistance and decreased fecundity are frequently reported in long-lived mutants with reduced activity of the insulin/IGF-1 signalling (IIS) pathway^29–31^. Therefore, our MetR condition recapitulates typical long-lived phenotypes.

Recently, it was reported that dietary cholesterol limitation is a strong modifier of lifespan extension by DR^32^. Since our holidic medium contained 0.1 g/L cholesterol, we increased its concentration to 0.3 g/L. Lifespan extension and the increased starvation resistance by MetR were still observed, supporting the conclusion that the present dietary regimen robustly increased the fly lifespan and starvation resistance independently of the cholesterol concentration (Supplementary Fig. 1c,d).

One of the characteristics of *Drosophila* ageing is a loss of climbing ability^33^. Measuring climbing ability is utilised as a way of measuring healthspan, and both DR and reduced IIS can improve ageing related climbing ability^34,35^. To quantify the climbing ability of individual flies, we developed a negative geotaxis assay (see Methods). The flies fed with the MetR diet for four weeks showed significantly improved climbing ability, although this phenotype was not observed in *w^Dah^*female flies (Supplementary Fig. 1e). Taken together, our data suggest that the life-long restriction of dietary Met recapitulates the benefits of protein restriction and thereby enhances the flies’ healthspan.

### Methionine restriction decreases Met metabolites

Met is a precursor of S-adenosylmethionine (SAM), a versatile methyl donor required for various methyltransferases (Fig. 1h). Upon methylation reaction, S-adenosylhomocysteine (SAH) is produced from SAM as a byproduct. SAH is further metabolised into homocysteine and then enters the transsulfuration pathway to make cysteine through cystathionine. On the other hand, free Met is converted into an oxidatively-damaged Met (methionine sulfoxide, MetSO). From LC‒MS/MS analysis of these metabolites, we confirmed that whole body Met levels were decreased under one week of MetR (Fig. 1i). All the detected Met metabolites were decreased (Fig.1j-m). The homocysteine level was below the detection limit in our analysis. Interestingly, the extent to which each metabolite decreased upon MetR largely varied. For instance, SAM and SAH levels were only decreased to half of the control level, when the Met level was only 6.34% (Fig. 1i-k). In contrast, the level of MetSO was sharply downregulated to 0.227% of the control upon MetR. This resulted in a 28-fold decrease in MetSO when compared to the decrease in internal Met, suggesting the presence of an active mechanism to downregulate MetSO levels during MetR. Considering that the absolute level of MetSO in control animals was ten times greater than that of Met, a large amount of Met was physiologically damaged (Fig. 1m).

We also noticed that other AA levels were affected during MetR. Threonine, asparagine, glutamine, and glycine were increased, and leucine, phenylalanine, tryptophan, and tyrosine were decreased (Supplementary Fig. 1f). Although we do not know the mechanisms by which other AAs are increased or decreased, methionine metabolism is coupled to these amino acids via one carbon metabolism or mitochondrial metabolism^36–38^.

### Early-life MetR extends lifespan

Next, we asked whether MetR in early or late adult life influences lifespan. For this experiment, the flies were fed with the MetR diet only for four weeks, and with the control holidic medium for the rest of their lives. Interestingly, feeding the MetR diet only for four weeks (day5-32) extended lifespan, almost as effectively as lifelong MetR, in female Canton-S (Fig. 2a). In contrast, flies fed with the control diet in the first four weeks did not live longer than non-restricted controls even though Met was restricted for the rest of their lives (Fig. 2a). This phenomenon was also reproduced in *w^iso^*^31^ fly strain using a slightly different MetR condition (1 mM Met in control vs 0.15 mM Met in MetR) (Fig. 2b, c), which was used in a previous report^19^. In this experiment, we restricted Met for the same duration (day5-day32 for early MetR and day32-day58 for late MetR) to compare its effects in young and old flies. Analysing the data using cox proportional hazards revealed that the timing of methionine restriction changed its effects on lifespan; early MetR extended lifespan relative to non-restricted controls, while MetR later in life did not (Fig. 2b, c; Supplementary Table 1). Interestingly, when we used the same protocol on *w^Dah^* females, both early and late MetR had similar effects in that they both extended lifespan, although early MetR again had a stronger effect on median lifespan (14.5% increase) than later life MetR (9.7% increase) (Fig 2.d, e; Supplementary Table 1). Importantly, upon returning MetR flies to the non-restricted control diet, the internal Met level rapidly recovered back to the control level in *w^is^*^o31^, negating the possibility that the Met level is irreversibly decreased by early MetR (Fig. 2f). Therefore, MetR in early life can robustly increase the lifespan of females with various genetic backgrounds and this effect is diminished or lost when MetR is applied later in life.

**Fig. 2.**
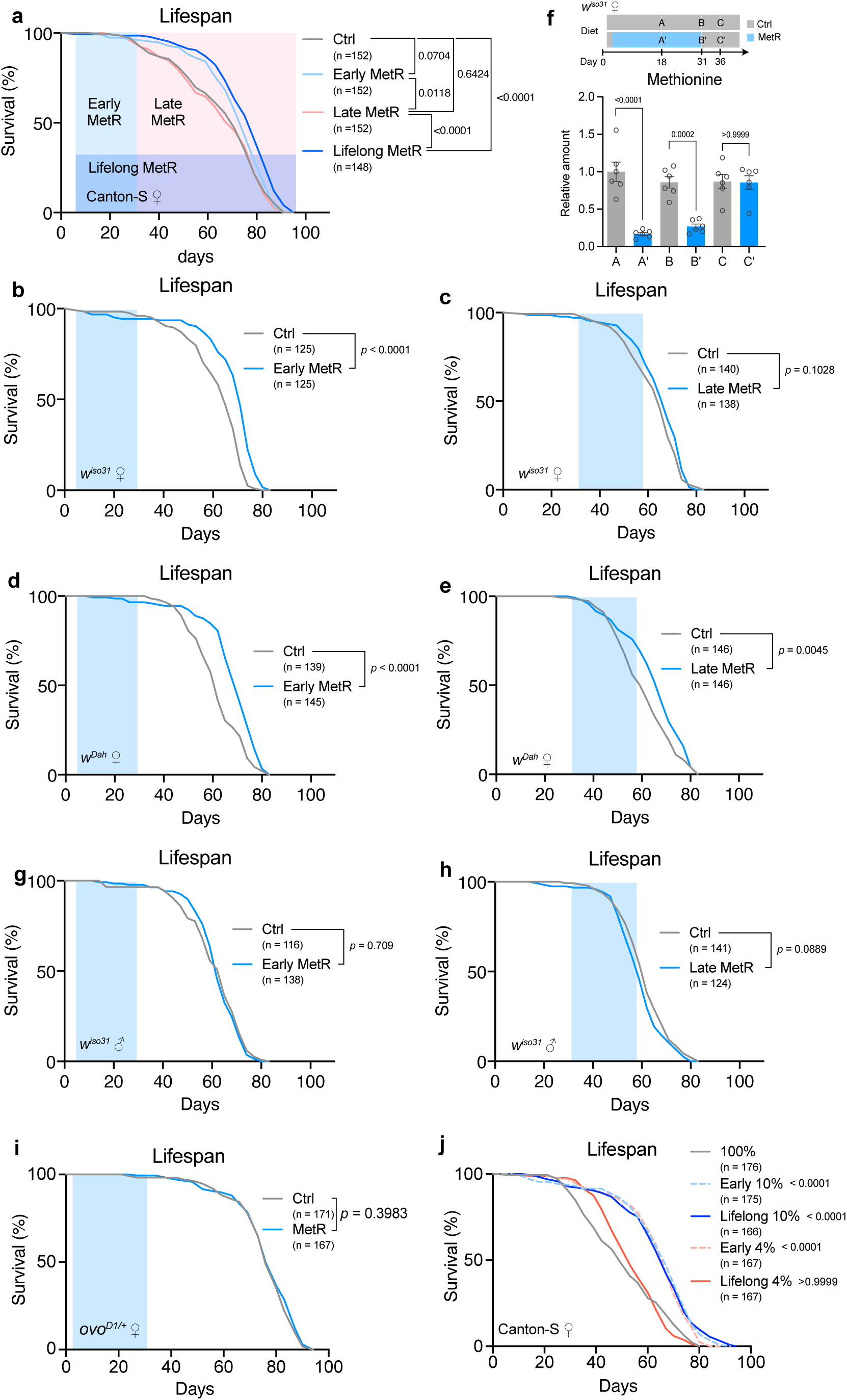
Early adult methionine restriction extends female lifespan. **a,** Lifespans of female Canton-S flies fed with or without a methionine-restricted diet during their early life (Day 5-31), late life (Day 31-), or whole life (Day 5-). Sample sizes (n) are shown in the figure. For the statistics, a log-rank test was used. **b-e,** Lifespans of female flies of *w^iso31^*(**b, c**) or *w^Dah^*(**d, e**) fed with or without a methionine-restricted diet in early life (**b, d**) or late life (**b, d**). Sample sizes (n) are shown in the figure. For the statistics, a log-rank test was used. **f,** Quantification of methionine upon its restriction over several time courses. n = 6. For the statistics, one-way ANOVA with Holm-Šídák’s multiple comparison test was used. **g, h,** Lifespans of male flies of *w^iso31^* fed with or without a methionine-restricted diet in early life (**f**) or late life (**g**). Sample sizes (n) are shown in the figure. For the statistics, a log-rank test was used. **i,** Lifespans of female *ovo^D1/+^* flies fed with or without a methionine-restricted diet during early life. Sample sizes (n) are shown in the figure. For the statistics, a log-rank test was used. **j,** Lifespans of female Canton-S flies fed a methionine-restricted (10% or 4%) diet compared to 100% AA during their early life or whole life. Sample sizes (n) are shown in the figure. For the statistics, a log-rank test was used. For the graph, the mean and SEM are shown. Data points indicate biological replicates.

In most cases, male flies do not exhibit large lifespan responses to dietary restriction. We did not observe any lifespan extension in response to early or late MetR in *w^iso^*^31^ males (Fig. 2g,h). We also tested the lifespan in *ovo^D1/+^* mutant female flies, in which no egg production was observed. The mutant had a relatively longer lifespan than fertile females in the control diet, and MetR in the first four weeks did not increase female lifespan further, suggesting that reproduction is indispensable for lifespan extension during early MetR (Fig. 2i).

A recent study has shown that overexpression of bacteria-derived methioninase in *Drosophila* can break down internal Met and increase lifespan without decreasing the level of other amino acids^39^. To assess if we observed extended lifespan because of reduced methionine or via an interaction between reduced methionine and other amino acids that we modified, we analysed lifespan of female Canton-S flies fed with holidic media containing 100% AA (containing 4 mM Met) with either 10% (0.4 mM Met) or 4% (0.16 mM Met, which is equivalent to 10% Met when 40% AA is used) of Met levels throughout life or only in early life. We found that both early and lifelong restriction of Met to 10% in 100% AA background can extend lifespan, but lifelong restriction of Met to 4% cannot (Fig. 2j). Intriguingly, limiting 4% Met restriction to early life only fully extended lifespan (Fig. 2j). This striking observation suggested that 1) decreasing all AA to 40% is not mandatory for MetR-longevity and 2) harsher MetR can extend lifespan when limited to early adulthood. Taken together, these data suggest that there might be a critical window for MetR to exert its maximal benefit and that early MetR likely induces a prolonged effect on the physiology.

### Transcriptomic response to dietary methionine declines in aged flies

Given that the later MetR has only a minor impact on fly lifespan, we speculated that aged flies become less responsive to the MetR diet. To determine how young and old flies react to dietary Met, we performed an RNAseq analysis using the female gut. We targeted the gut for this analysis because previous studies have demonstrated it to be critical for lifespan responses to DR^28,40^. To maximise the response, we completely removed Met (Met-) from the diet and compared it with the complete holidic medium, which included the original amounts of all amino acids. Principal component analysis (PCA), quantification of differentially expressed genes (DEGs, FDR<0.01, fold change>1.5), and heatmap analysis clearly demonstrated that the Met- diet induced a strong transcriptional shift in the young gut that was much blunted in the aged gut (Fig. 3a-c, Supplementary Data 1). In the PCA, the PC1 axis separates the animals by age, which was slightly shifted towards the left by just one day of feeding of the Met- diet in young flies, but not in aged flies (Fig. 3a). This suggested that the Met- diet can acutely shift the transcriptome towards rejuvenating the physiological age of the flies. It is also clear, that dietary treatment clearly separates the samples along the secondary PC axis in young flies, but this separation is almost eliminated in aged flies.

**Fig. 3.**
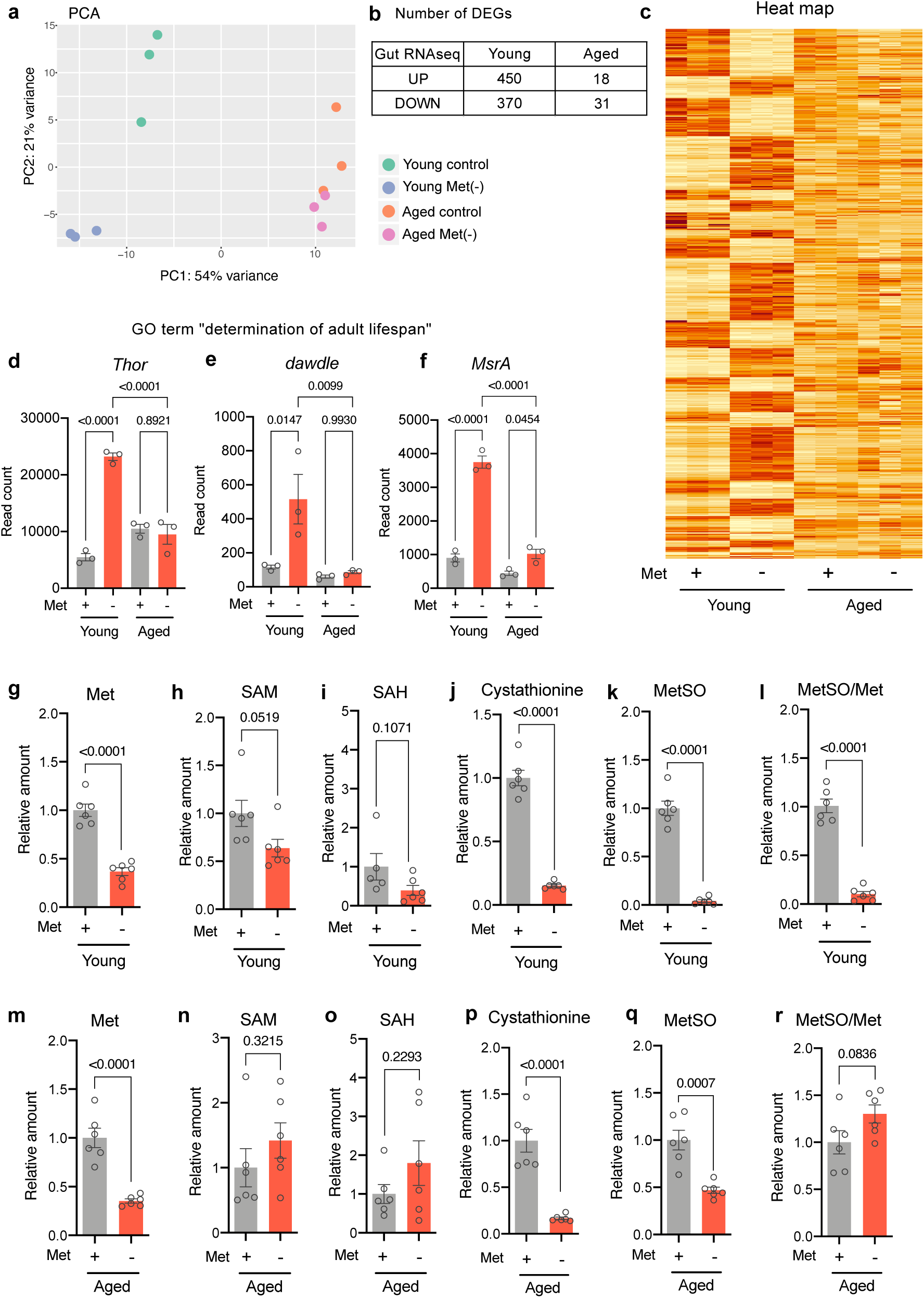
A blunt transcriptomic response to short-term methionine depletion in old flies. **a-c** Principal component analysis (**a**), the number of differentially expressed genes (**b**), or heatmap analysis (**c**) of RNAseq analysis of intestines in young and aged female *w^iso31^* flies upon methionine depletion for 24 hours. **d-f,** Read counts of genes categorised as the Gene Ontology of “determination of adult lifespan” in the RNAseq analysis of intestines in young and aged female *w^iso31^* flies upon methionine depletion for 24 hours. For the statistics, one-way ANOVA with Holm-Šídák’s multiple comparison test was used. **g-r,** Quantification of methionine metabolites and the oxidative product upon methionine depletion for 24 hours in young (**g-l**) or aged (**m-r**) female *w^iso31^* flies. n = 6. For the statistics, a two-tailed Student’s *t* test was used. For all graphs, the mean and SEM are shown. Data points indicate biological replicates.

We found 820 DEGs upon 24-hour feeding of the Met- diet in the young female gut. In clear contrast, we found only 49 DEGs in the guts of aged females fed the same Met- diet, indicating a 94% reduction in the number of DEGs during ageing (Fig. 3b). Gene Ontology analysis revealed that the short-term Met depletion changed gene sets related to “determination of adult lifespan”. Several genes included in the GO term were induced by early, but not late, Met depletion (Fig. 3d-f, Supplementary Fig. 2a). These data suggest that the aged gut was much less responsive to the lack of dietary Met. The upregulated genes included *Thor/4E-BP*, a negative regulator of energy-consuming protein synthesis, which is a common target of many nutrient-responsive pathways. Increased *Thor* is a hallmark of the starvation response, suggesting that the tissue sensed the lack of nutrients in the Met- diet, but the older gut did not. Other starvation-related processes, such as autophagy, stress response, lipid metabolism, nutrient transport, and mTOR inactivation were also enhanced in young but not older guts (Supplementary Fig. 2b-k).

One simple possible reason for the blunted response to MetR in the aged flies is that their internal Met levels might not have been decreased to the same extent as in young flies. To test this hypothesis, we measured Met and its metabolites by LC‒MS/MS analysis (Fig. 3g-r). The whole-body Met levels in both young and aged flies were decreased by 24-hour feeding with the Met- diet (Fig. 3g, m). We noticed that many Met metabolites, such as SAM, SAH, and MetSO, were decreased by the Met- diet in young flies (Fig. 3h,I,k,l), but the effects were mild in the aged flies (Fig. 3n,o,q,r). Nonetheless, the level of cystathionine was similarly decreased even in the aged flies (Fig. 3j,p). The reason why cystathionine can be decreased without changing SAM and SAH levels is unknown, but one can assume that another branch of Met metabolism, such as the Met salvage pathway, in which SAM is converted into polyamines, could be inhibited in the aged flies. In any case, this observation negates the possibility that all Met metabolism was simply slowed down during ageing. Taken together, these results indicate that in aged flies, MetR decreases the levels of internal Met and some Met metabolites but provokes only a partial transcriptomic response.

### Transcriptome analysis of the gut and the fat body to the MetR diet

Complete depletion of dietary Met resulted in a large shift in the transcriptome (Fig. 3a-c). To understand the transcriptional response to early MetR, we performed a time-series ‘‘mRNAseq analysis in the gut at six hours, two weeks, and four weeks of the lifespan-extending MetR diet. Female *w^Dah^* flies fed a 0.15 mM Met diet were compared with the control flies fed with 1 mM Met. The results showed that approximately 30 genes were differentially expressed under MetR conditions at six hours or two weeks (Fig. 4a, Supplementary Data 2). In contrast, four weeks of MetR increased the number of DEGs to as much as 252 (Fig. 4a). There was no gene commonly changed at all three timepoints. Eighteen upregulated genes were shared by two week- and four week-MetR, while one (six hours vs four weeks) or two (two weeks vs four weeks) downregulated genes were common (Fig. 4b-f). Amongst these genes in common to more than one timepoint, the upregulated genes included *methionine sulfoxide reductase A* (*MsrA*, also known as *Eip71CD*), transporters (CG3036 and CG8785), genes related to lipid metabolic processes (*Lipid storage droplet-2* (*Lsd-2*) and *Acetyl Coenzyme A synthase* (*AcCoAs*)), and serine proteases (CG18180 and *Jonah 99Fii* (*Jon99Fii*)) (Fig. 4d). Both *MsrA* and *Lsd-2* were also upregulated under the Met- conditions (Fig. 3f, Supplementary Fig. 2e), suggesting that these genes are robustly responsive to dietary Met. The downregulated genes were *Glutathione S transferase E7* (*GstE7*), *CCHamide-2* (*CCHa2*), and an unknown gene CG3902 (Fig. 4e,f). CCHa2 is an appetite-regulating peptide that is known to promote systemic insulin signalling^41^. Thus, decreased expression of *CCHa2* may lead to a pro-longevity state triggered by reduced IIS activity. Interestingly, we found from i-*cis*Target^42^ that many genes containing the foxo consensus binding site were induced in the gut upon four weeks of MetR, further indicating induction of IIS (Fig. 4g).

**Fig. 4.**
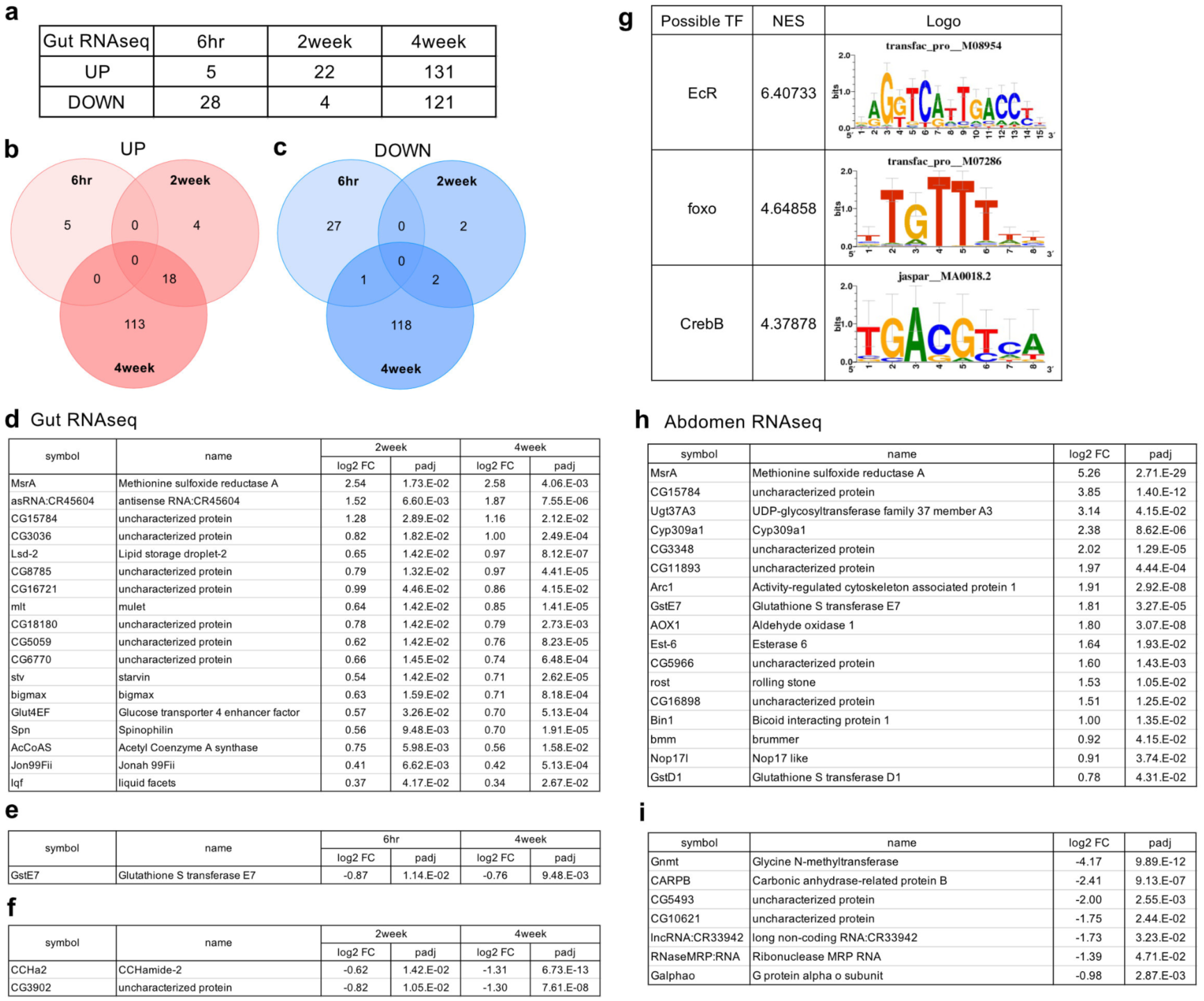
Time-course transcriptomic responses to methionine restriction. **a-c,** Numbers (**a**) and Venn diagrams (**b, c**) of differentially expressed genes of the 3’ RNAseq results from the intestines of female flies upon acute (six hours) or chronic (two or four weeks) methionine restriction. **d,** List of commonly upregulated genes in the female intestines after two and four weeks of methionine restriction. **e,** Commonly downregulated genes in the female intestines after six hours and four weeks of methionine restriction. **f,** Commonly downregulated genes in the female intestines after two and four weeks of methionine restriction. **g,** Possible transcription factors related to the transcriptional responses upon methionine restriction analysed by i-*cis*Target. **h, i,** Lists of commonly upregulated (**h**) and downregulated (**i**) genes in the female abdomens upon four weeks of methionine restriction.

To elucidate the systemic response to the MetR diet, we also performed RNAseq analysis of the abdomen upon four-week MetR, which contains the fat body, the major metabolic organ in *Drosophila*. The digestive tract and reproductive organs were carefully removed for this analysis. Strikingly, the most upregulated gene in the abdomen was *MsrA* (Fig. 4h) as was the case in the gut (Fig. 4d). We also noticed that a lipase *brummer* (*bmm*) was upregulated in the tissue, which implied altered lipid metabolism in the flies fed the MetR diet for four weeks. *bmm* is another foxo target, suggesting that foxo is activated in both the gut and the abdomen^43,44^.

In contrast, *glycine N-methyltransferase* (*Gnmt*) was strongly suppressed in the abdomen upon MetR (Fig. 4i). Gnmt is an enzyme that produces sarcosine by utilising SAM, and thereby regulates the SAM levels^45^. The phenomenon that *Gnmt* expression is suppressed by low methionine intake was observed consistently in our previous study^21^. This SAM buffering system contributes to maintaining SAM levels by avoiding excess SAM utilisation during MetR (Fig. 1j, Fig. 3h). Considering that inhibiting *SAM synthase* (*sams*) also results in a decrease in *Gnmt* expression^46^, there must be a system to downregulate *Gnmt* mRNA upon detection of decreased SAM levels. The mechanism of this SAM-dependent regulation of *Gnmt* remains to be elucidated.

### MetR induces *MsrA* in a foxo-dependent manner

Given that *MsrA* was the most upregulated gene by four weeks of MetR in both the abdomen and the gut, we focused on its regulatory mechanism and function. Msr is an evolutionarily-conserved antioxidant enzyme required for repairing oxidatively-damaged Met (MetSO) back into functional Met^47,48^. It reduces both free and protein-based Met-*S*-SO. In physiological conditions, free-MetSO exists in hemolymph at a level four times more than Met (Supplementary Fig. 3a). This implies that Met is oxidized in body fluids, or MetSO formed in tissues is circulated throughout the body. It has already been shown that overexpression of *MsrA* can confer oxidative stress resistance and extend lifespan in *Drosophila*^49^. Ectopic expression of a yeast *Msr,* which can reduce free Met-*R*-SO, extends the *Drosophila* lifespan, especially under a Met-rich diet^50^. From these previous reports, we speculated that induction of *MsrA* is one of the functional genes mediating the healthspan-extending effect of MetR.

We first confirmed by quantitative RT–PCR that the induction of *MsrA* was robustly observed when using a lifespan extending MetR regimen (i.e., 1.6 mM for Control vs. 0.16 mM for MetR in *w^Dah^*, Fig. 5a-c). *MsrA* could be upregulated as early as three days of MetR in both the gut and abdomen of males and females (Supplementary Fig. 3b-e). Given MetR-longevity is marginal in males, the MetSO levels may not be the limiting factor for male lifespan. The *ovo^D1/+^* mutant females did not significantly increase *MsrA* expression upon MetR (Supplementary Fig. 3f), suggesting that *MsrA* induction in females depends on reproductive capacity. Interestingly, *MsrA* induction by four-week MetR was long-lasting at least two weeks after shifting back to the control diet, although the magnitude of induction decreased (Fig. 5b,c). Notably, the induction of *MsrA* was attenuated during ageing (Fig. 3f). In addition, MetSO was massively decreased by the Met- diet in young animals, whereas it was mild in aged ones (Fig. 3k,l,q,r). Therefore, the increased expression of *MsrA* gene and the concomitant decrease of MetSO, in both absolute levels and relative to the level of Met, by MetR was much greater in the young flies, which may account for the lack of benefit of MetR at the later age.

**Fig. 5.**
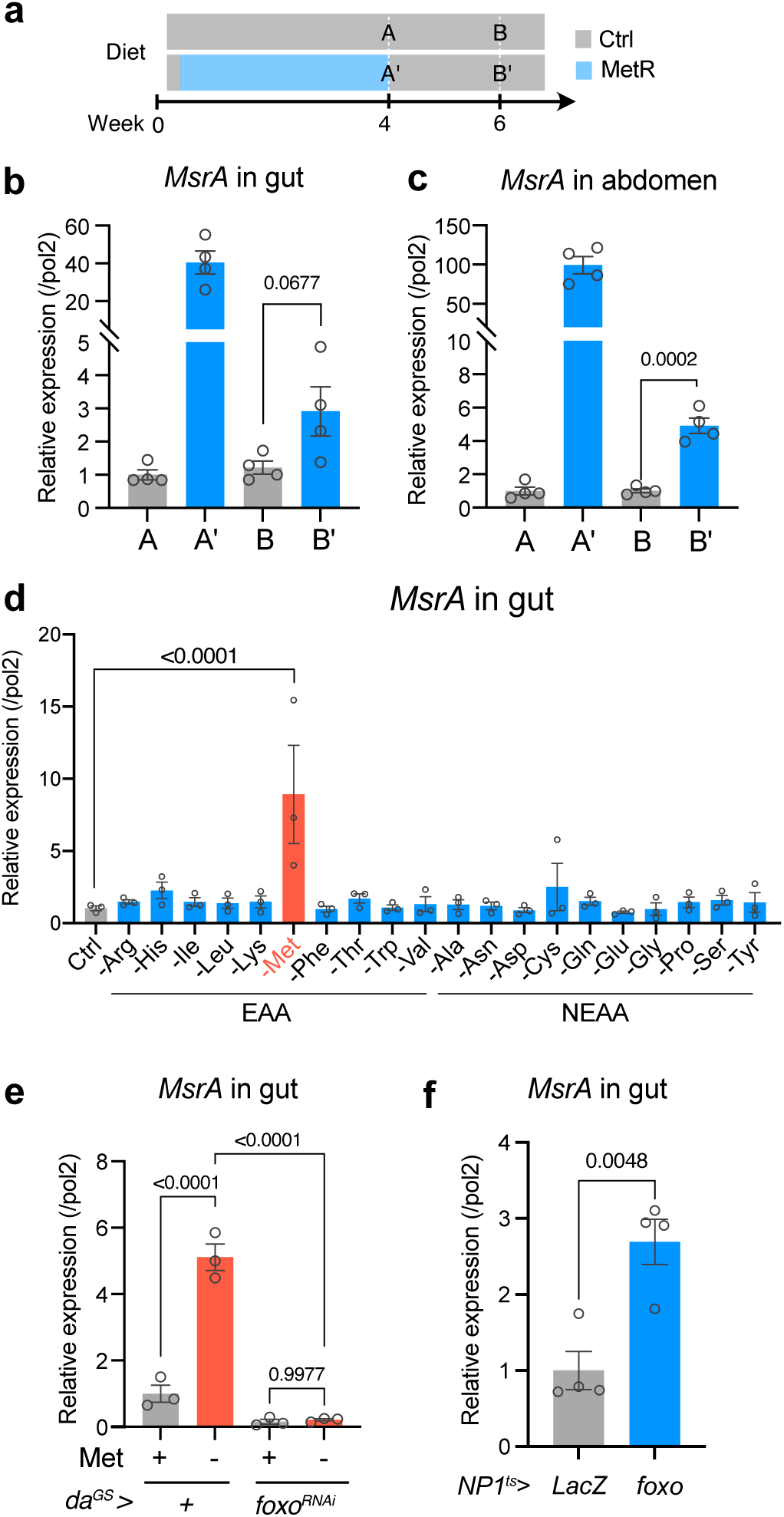
*MsrA* is induced by methionine restriction via foxo. **a-c,** Time course (**a**) and the results (**b, c**) of the quantitative RT‒PCR analysis of *MsrA* in the gut (**b**) and abdomen (**c**). For the statistics, a two-tailed Student’s *t* test was used. **d,** Quantitative RT‒PCR analysis of *MsrA* in the guts fed with single amino acid-restricted diets for 24 hours. For the statistics, one-way ANOVA with Dunnett’s multiple comparison test was used. **e,** Quantitative RT‒PCR analysis of *MsrA* in the guts fed a methionine-restricted diet for 24 hours. Knockdown of *foxo* using the drug-inducible whole-body driver *da^GS^*. RU486 (20 μM) was added to the medium. For the statistics, one-way ANOVA with Holm-Šídák’s multiple comparison test was used. **f,** Quantitative RT‒PCR analysis of *MsrA* in the gut. Overexpression of *foxo* for four days using an enterocyte driver *NP1-Gal4* with temperature sensitive *tub-Gal80^ts^*. The flies were fed a standard yeast-based diet. For the statistics, a two-tailed Student’s *t* test was used. For all graphs, the mean and SEM are shown. Data points indicate biological replicates.

Interestingly, depletion of no other single AA from the diet, except for Met, for 24 hours did not increase *MsrA* expression in the gut (Fig. 5d), suggesting that *MsrA* is specifically regulated by dietary Met. *MsrA* is a known target of the pro-longevity transcription factor Daf-16 in *C. elegans*, foxo in *D. melanogaster*, and FOXO3a in a cultured human cell line^49,51^. Considering that foxo was predicted to be a mediator of the MetR-responsive transcriptional shift (Fig. 4g), we hypothesised that MetR increases *MsrA* through the activation of foxo. Consistent with this, *MsrA* induction by the Met- diet was blocked by *foxo*-RNAi (Fig. 5e). Furthermore, *foxo* overexpression in the adult gut for four days was sufficient for *MsrA* induction (Fig. 5f). These data indicate that foxo induces *MsrA* during MetR.

### MsrA function is required for MetR-induced longevity

To test whether MsrA is functionally relevant to MetR-induced lifespan extension, we asked if loss of function of *MsrA* alters lifespan. A mutant fly line *MsrA^EY^*^05753^ with a P-element insertion in the second exon of the *MsrA* locus was utilised (Fig. 6a). *MsrA^EY^*should be a null allele, as it has a large insertion in the coding sequence common to all isoforms. We confirmed by qRT–PCR that there was no detectable level of *MsrA* transcripts in the mutant (Supplementary Fig. 4a). We backcrossed the *MsrA^EY^*^05753^ line into *w^Dah^* to minimise the effect of genetic background on lifespan and metabolic analyses. The mutant was viable and fertile and did not have a shortened lifespan on the control diet (Fig. 6b). This is consistent with a previous report, demonstrating the redundant functions of MsrA and MsrB^52^. Surprisingly, the lifespan of the *MsrA^EY^* female fly is longer than that of *w^Dah^* on the control diet (Fig. 6b). This could be because there is an increase in mild oxidative stress in the mutant flies that induces a13ermeticc effect, reinforcing the stress resistance and extending lifespan. Indeed, the mutant showed an increased resistance to hydrogen peroxide (Supplementary Fig. 4b). Strikingly, the *MsrA^EY^*mutant did not show lifespan extension upon MetR (Fig. 6b, Supplementary Table 2). Note that, in this analysis, *MsrA^EY^* mutant flies have an insertion of mini *white* gene, while the control flies do not. Thus, while the dose of *white* could have affected the lifespan, although it cannot account for the change in response to MetR since both red and white-eyed flies have extended lifespan in response to MetR (Fig. 2a, b, d).

**Fig. 6.**
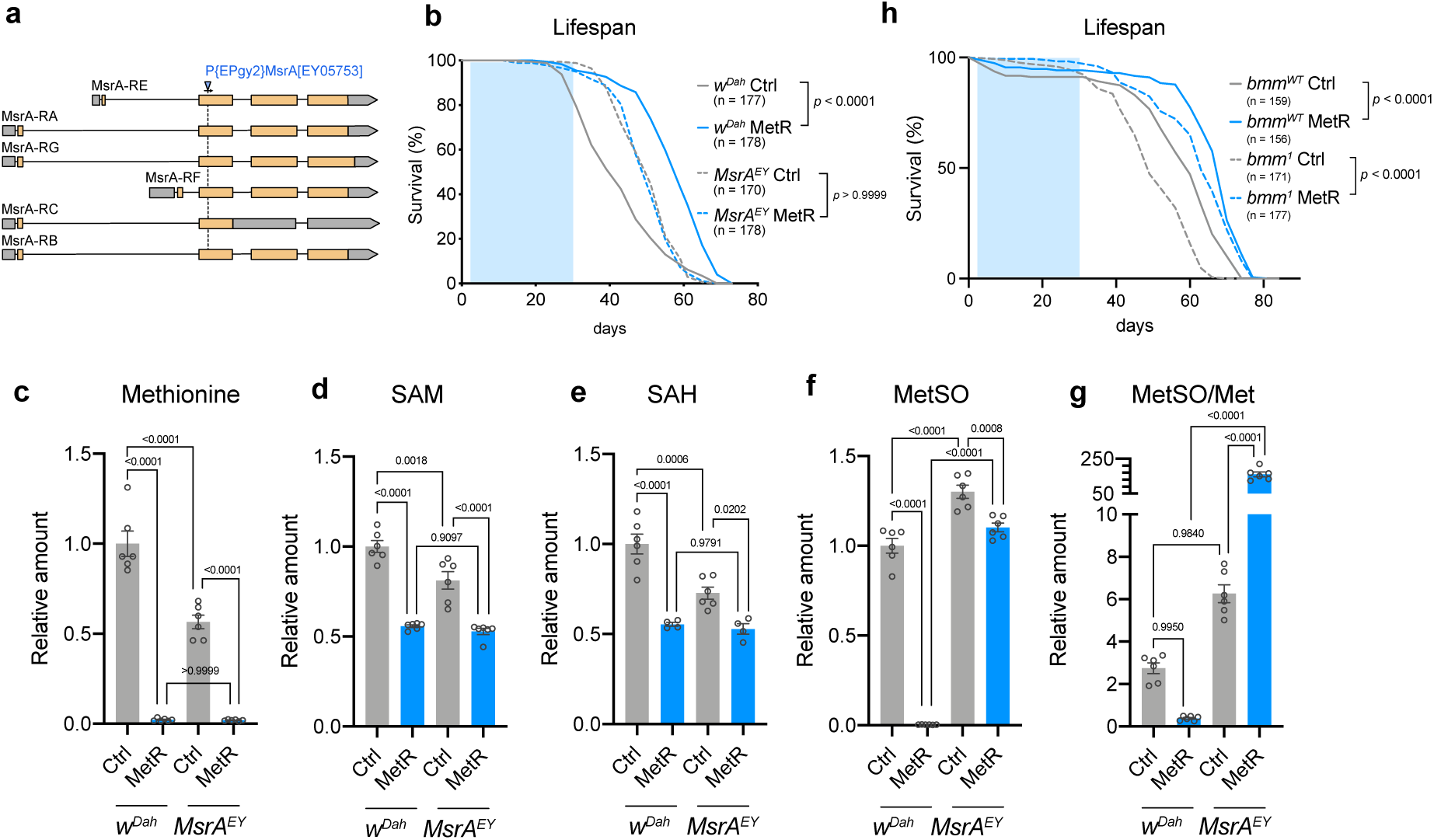
Loss of *MsrA* abolishes lifespan extension by methionine restriction. **a,** The gene structure of *MsrA* and the insertion site of the p-element in the *MsrA^EY^* mutant. **b,** Lifespans of female flies of *w^Dah^* and *MsrA^EY^* fed with or without a methionine-restricted diet in early life. Sample sizes (n) are shown in the figure. For the statistics, a log-rank test was used. **c-g,** Quantification of methionine metabolites and the oxidative product in female *w^Dah^* or *MsrA^EY^* flies upon methionine restriction. n = 6. For the statistics, one-way ANOVA with Holm-Šídák’s multiple comparison test was used. **h,** Lifespans of female flies of *bmm^WT^* and *bmm^1^* fed with or without a methionine-restricted diet in early life. Sample sizes (n) are shown in the figure. For the statistics, a log-rank test was used. For all graphs, the mean and SEM are shown. Data points indicate biological replicates.

In *MsrA^EY^* mutants fed with the control diet, we found that Met/SAM/SAH was decreased while MetSO was increased (Fig. 6c-g). These data suggest that up to half of the free Met is constantly recovered from MetSO under physiological conditions. As observed in Canton-S, all the Met metabolites in *w^Dah^* were decreased by one-week-MetR, among which the decrease in the level of MetSO was the strongest (Fig. 1i-m, Fig. 6c-g). This led to a decrease in the MetSO/Met ratio, which was also the case in the Met- diet in young *w^Dah^* flies (Fig. 1m, 3l, 6g). As anticipated, while Met, SAM, and SAH were still decreased, the MetSO reduction was blunted and the MetSO/Met ratio was increased by MetR in the *MsrA^EY^* mutant (Fig. 6g). These data are similar to what we observed when we subjected aged flies to Met depletion (Fig. 3q, r). Together, these data indicate that the MetR-induced decrease of MetSO requires MsrA in a manner that is consistent with it having a causal role in lifespan extension.

### Lipid metabolism and the transsulfuration pathway may not be involved in MetR- induced longevity

MetR promoted lipid accumulation in the gut of wild-type flies (Supplementary Fig. 1a, b) and upregulated the gene expression of *Lsd-2* and *bmm* (Fig. 4d,h, Supplementary Fig. 2e). This phenotype implies that MetR promotes lipid turnover, phenocopying what happened during conventional DR^28,53^. To test whether altered lipid metabolism contributes to early MetR-longevity, we analysed a *bmm^1^* mutant and its control animals with the same genetic background^43,54^. In this mutant, neutral lipid in the gut was already abundant under the control diet, and were mildly upregulated upon MetR, which correlated well with the flies’ enhanced starvation resistance (Supplementary Fig. 5a,b). Mutation of the brummer lipase did not compromise the lifespan extension by MetR (Fig. 6h, Supplementary Table 3). Although we cannot rule out the possibility that blocking other lipid metabolic genes or upstream transcription factors could alter the MetR longevity, these data suggested that the altered lipid metabolism contributed little to the lifespan regulation in this context.

Methionine metabolism is coupled with cysteine metabolism through the transsulfuration pathway (TSP) (Supplementary Fig. 6a). To test whether TSP is related to MetR-longevity, we used the TSP inhibitor propargylglycine (PPG). First, we fed female flies with various concentrations of PPG during MetR and quantified the internal Met metabolites. As expected, cystathionine accumulated in response to PPG in a dose dependent manner, while the levels of Met, SAM, SAH and MetSO did not respond to the PPG concentration (Supplementary Fig. 6b-f). The administration of PPG shortened the female lifespan (Supplementary Fig. 6g). Strikingly, however, we still observed lifespan extension by early-adult MetR (Supplementary Fig. 6g). From these data, we concluded that TSP is not involved in the MetR-induced longevity.

### Single-cell RNAseq analysis of the gut upon methionine restriction

Lastly, we tried to describe how early MetR influences tissue homeostasis and thereby maintains healthspan by single-cell RNAseq analysis of the female gut. The midgut samples were dissected from *w^Dah^* flies in young (day6), aged (day40, 1.6 mM Met for five weeks), and aged-MetR (day40, 0.16 mM Met for five weeks) conditions (Fig.7a). The Malpighian tubules, the hindgut, and the proventriculus were removed manually. The estimated numbers of sequenced cells for young, aged, and aged-MetR were 10364, 15604, and 20313, respectively.

After batch correction and data cleaning, the cells were clustered into 24 different cell types expressing specific genes, which were visualised using a uniform manifold approximation and projection (UMAP) plot (Fig. 7b, Supplementary Fig. 7a-c). As commonly observed in the fly gut, we found four major cell types: intestinal stem cell (ISC), enteroblast (EB), enterocyte (EC), and enteroendocrine cell (EE). The number of cell clusters in each cell type varies among studies^55–57^. In our case, we observed seven progenitor (ISC/EB) clusters, nine EC clusters, and seven EE clusters. During ageing, the proportions of cell numbers in some clusters changed. For example, there were decreases in several progenitors (EB/ISC, EB1, and EB2), and increases in some enterocyte clusters (Fig. 7c-e). Among these changes, MetR restored the increased proportions of clusters of posterior enterocytes (pEC1-pEC4) to more youthful levels (Fig. 7d). These results suggest that one of the benefits of MetR can be the suppression of the (relative) expansion of posterior enterocytes.

**Fig. 7.**
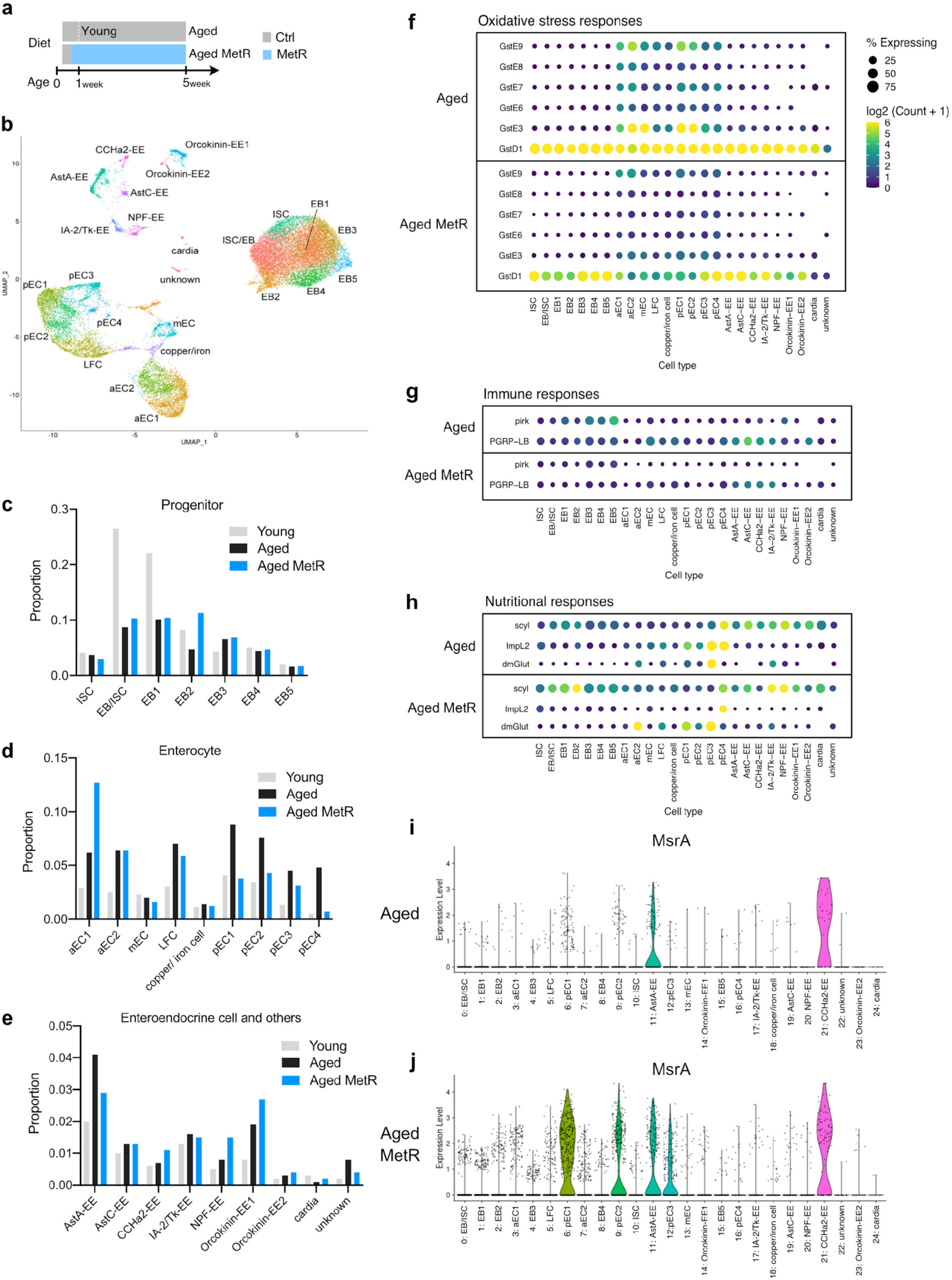
Single-cell RNAseq analysis of the guts of methionine-restricted flies. **a,** Experimental time course of gut sampling for single-cell RNAseq analysis upon methionine restriction. **b,** UMAP plot and annotated cell types from the single-cell RNAseq analysis. **c-e,** Proportions of cell types in progenitor cells (**c**), enterocytes (**d**), and enteroendocrine cells (**e**). **f-h,** Dot plots of genes related to immune responses (**f**), oxidative stress responses (**g**), or nutritional responses (**h**). **i, j,** Violin plots of *MsrA* expression in the aged gut (**i**) or the aged gut with methionine restriction (**j**).

Next, we investigated what kind of transcriptomic change occurs in each cell cluster (Supplementary Data 3). A heatmap analysis of the DEGs in each cell type showed a rough trend in which age-related shifts in the transcriptional signature were recovered back by MetR, especially in progenitors and enterocytes (Supplementary Fig. 8a,b). In contrast, MetR induced a distinctive transcriptome pattern in enteroendocrine cells (Supplementary Fig. 8c). PCA of the data also supported this observation (Supplementary Fig. 9). For instance, in progenitors, we observed a shift to the left in the PC1 axis by ageing, which was shifted back to the right by MetR (Supplementary Fig. 9a). This was also the case in anterior and posterior enterocytes, where the age-dependent shift along the PC1 axis was recovered under MetR condition (Supplementary Fig. 9b,d). However, the degree of the “rescue” of the age-related transcriptomic shifts varied greatly among clusters.

It has been observed that the number of damaged/active ISCs rises in the aged gut^57^. However, in our analysis, we did not observe this rise, but rather decreased numbers of some progenitor cells (Fig. 7c). One possible reason for this is that our flies were fed with the holidic medium, which contains more preservatives. This diet generally resulted in lower numbers of gut bacteria, which are known to mediate the age-related increase in ISC expansion^58^. Another possibility is that the data cleaning process removed “unordinary” cells in the aged samples from the clustering analysis, due to the disturbed gene expression signature. For the Aged sample, only 31.7% of the sequenced cells were clustered (4946/15604), whereas this was 99.5% for the Young sample (10310/10364). For the Aged-MetR sample, this number was increased to 48.6% (9863/20313), implying that their transcriptional profile was more youthful, perhaps indicating delayed tissue ageing.

Considering that there were also large differences in the quality of the data (RNA volume, sequence reads, number of genes) between the Young and the two Aged samples, it seems difficult to fairly compare the absolute gene expression between the two ages. Thus, we carefully looked at DEGs that were changed between Aged and Aged-MetR. We noticed that some DEGs were commonly changed among cell clusters. Several GSTs were decreased by MetR (Fig. 7f), which was consistent with the bulk RNAseq analysis (Fig. 4e). Decreases in GSTs expression imply a lower level of oxidative stress in the tissue. It was also clear that expression of the immune-related genes *pirk* and *PGRP-LB* were decreased by MetR, suggesting attenuation of the “inflammatory” state in the aged gut (Fig. 7g). Recent scRNAseq analysis in the fly gut also found that suppressing an innate immune Imd pathway activity in ISCs autonomously attenuates age-related ISC overproliferation^59^. It was also observed that some nutritional response genes were affected (Fig. 7h). Notably, *MsrA* was induced in many cell types, especially in the posterior enterocyte clusters (Fig. 7i,j). This was correlated with the MetR-driven decrease in the number of cells and GSTs expression in the posterior enterocytes (Fig. 7d, f). Taken together, single-cell RNAseq analysis of the gut suggested that MetR delayed the tissue ageing signature. It also produced a resource of many addressable hypotheses to understand how a nutrient impacts the health of each cell type in the tissue.

## Discussion

There is an urgent need to develop anti-ageing interventions. Studies using model organisms have contributed to understanding the biology of ageing, which will eventually lead to the development of various methods of lifespan extension^60^. DR is thus far one of the most robust and practical applications for human society. However, restricting diet throughout life is not entirely appropriate since the effect of diet depends on the conditions, such as age^5^. In this study, we demonstrated that MetR in early adulthood efficiently extends fly lifespan, whereas that in later adulthood has a milder effect (Fig. 8). This may be at least partly because aged flies cannot trigger the beneficial transcriptional shift in response to the dietary manipulation, despite internal Met level being decreased, suggesting that ageing blunts the transcriptional response to a particular nutrient shortage. Our data imply the existence of a “methionine memory”, which can be defined by the amount of dietary Met to which the animal is exposed in its early life stage. They also suggested that MetR can help maintain the younger state, but once the anima’’s physiological condition goes beyond a threshold (middle to aged), it may not preserve the animal anymore.

**Fig. 8.**
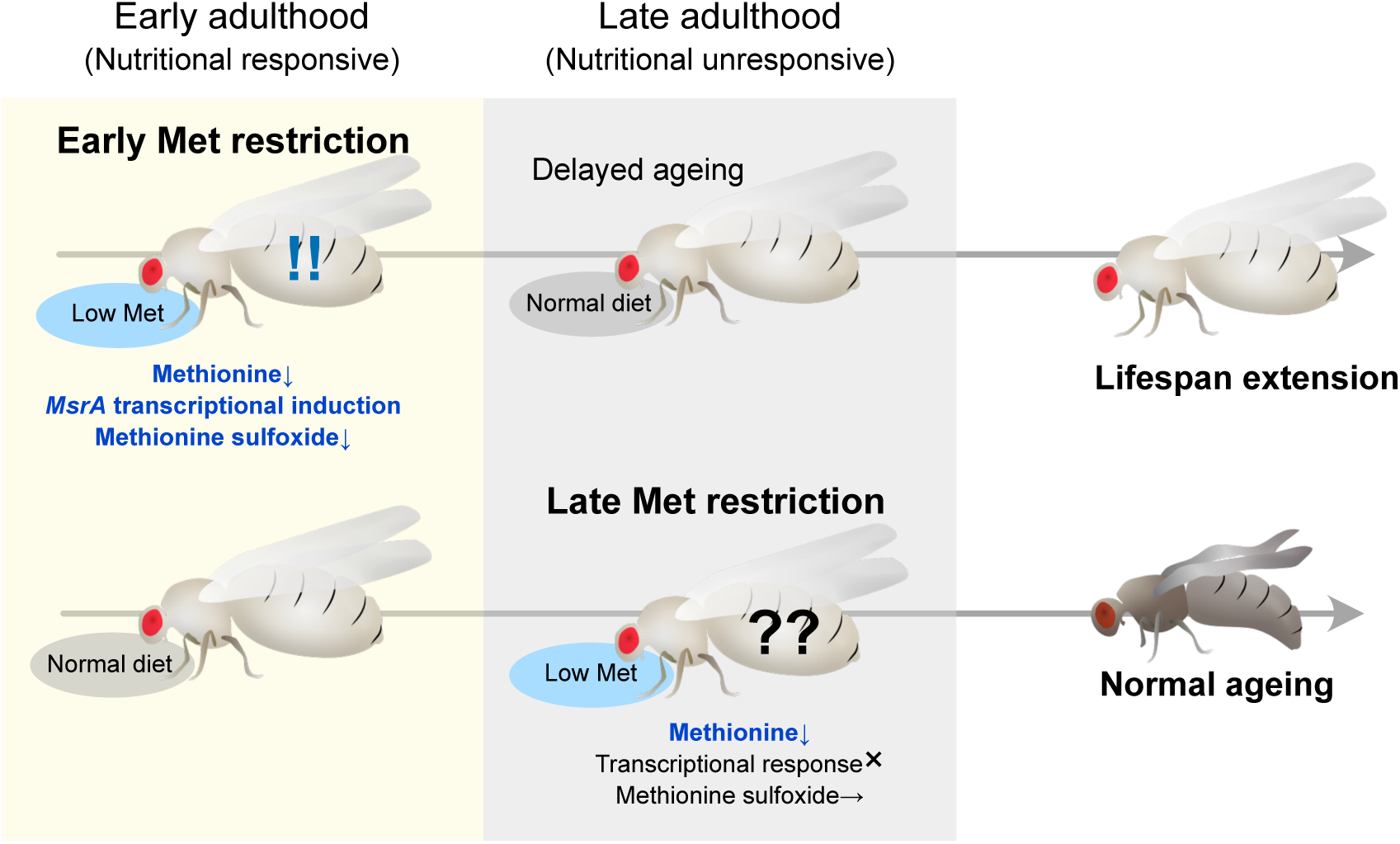
A model.

How MetR provokes the (prolonged) transcriptional response and lifespan extension is not entirely understood. Theoretically, decreased Met can be sensed through amino acid-sensing kinases such as general control nonrepressible 2 (GCN2) and mechanistic target of rapamycin (mTOR). Activation of mTOR in the fat body by nutrients can stimulate the secretion of dilps, which in turn activates insulin/IGF-1 like signalling (IIS) systemically^61^. This nutritional relay has been extensively studied in the larval stage, but the mechanism is also thought to be conserved in adults^62–64^. GCN2 activation leads to a transcription through activation transcription factor 4 (ATF4), while downregulation of the TOR/IIS pathway activates the transcription factors, REPTOR and foxo^65^. We found that foxo, ecdysone receptor (EcR), and CREB were predicted to be transcription factors involved in the MetR-induced transcriptomic response. Interestingly, both EcR and CREB transcription factors are known to be cofactors of ATF4. Thus, it is likely that these nutrient sensing machineries were all affected by MetR.

In this study, we found that *MsrA* was likely to be regulated by foxo. Increased *MsrA* expression two weeks post MetR suggests prolonged foxo activation. It is possible that decreased IIS activates foxo under MetR. However, at least in worms, *MsrA* overexpression enhances the nuclear localisation of DAF-16/foxo in the gut. Therefore, a foxo-MsrA feedforward loop can also mediate the effect of MetR. Careful analysis of foxo activation, including protein modification and localisation, is necessary to further understand the mechanistic basis. Additionally, it is intriguing to understand why the foxo-MsrA axis is specifically sensitive to Met, but not other aAs.

There is a general concept that ageing decelerates metabolism. However, Met depletion in both young and aged flies resulted in decreased levels of cystathionine, a downstream metabolite produced through TSP. Nonetheless, we observed increased SAM and SAH levels in aged flies upon Met depletion. Given that SAM seems to be a major regulator of intestinal homeostasis and organismal lifespan^21,66^, a lack of SAM decrease in aged flies would result in a blunted response to Met restriction. SAM is also a precursor for polyamines, which are reported to be mediators of DR-longevity in *Drosophila*^23^. If SAM is a major regulator of the foxo-MsrA axis, we could explain the Met specificity for the regulation of *MsrA* expression. This possibility needs to be carefully analysed in the future.

Our data suggested that *MsrA* is a functional target of MetR that drives the beneficial program that counteracts organismal ageing. It is reported in the genetic model of MetR that, the level of MetSO is also strongly decreased, although the direct contribution of *MsrA* to this phenotype was not tested^39^. The literature has already demonstrated that MsrA contributes to reducing MetSO and alleviating oxidatively damaged proteins and free non-functional Met. Interestingly, dimethyl sulfide (DMS) is reported to extend lifespan in *C. elegans* and in *Drosophila* in an *MsrA*-dependent manner^67^. In contrast, MsrA was reported to be nonessential for metabolic benefit of MetR in mice^68^. Whether MsrA is required for lifespan extension in other contexts in various organisms including mammals needs to be tested in the future.

This study did not identify the tissue where 1) Met is sensed and 2) *MsrA* functions to drive a pro-longevity mechanism. Previous research using genetic MetR models in *Drosophila* has shown that lifespan extension occurs when Met levels are decreased either in the gut or in the fat body (Parkhitko et al., PNAS 118, e2110387118, 2021). Their data suggest that MetR in at least these two tissues contribute to the organismal longevity. Another report indicates that overexpression of *gnmt* in the fat body has been found to extend lifespan (Tain et al., Aging, Cell, 2019). These results suggest that a decrease in Met/SAM levels in the gut or fat body can be a key factor in lifespan extension. This is consistent with the fact that we found *MsrA* to be induced in these organs during MetR. However, we also found that Met/SAM/MetSO are metabolites that can be transported through circulation (as they exist in hemolymph). Thus, changes in MsrA expression in multiple peripheral tissues, not only in a specific tissue, may be important to decrease systemic MetSO levels to an extent that extends lifespan.

It is possible that *MsrA* induction during MetR is an adaptation of cells to counteract Met shortage; perhaps MetSO can serve as a store of Met. Indeed, the absolute amount of MetSO is ten times higher than the free Met in female flies. Interestingly, recent work has shown that egg production in female *Drosophila* decays quickly (within three days) in response to deprivation for any one of eight of the essential amino acids^69^. By contrast, exposure to food lacking either histidine or methionine results in a much slower decay in egg production (seven days) indicating that the flies have internal reserves of methionine that they can call on to buffer against environmental fluctuations. This response may have evolved specifically for these essential amino acids because they appear to be consistently limiting for flies feeding on their natural diet of yeast^18,26,70^.

Once we know the mechanism of MetR-induced *MsrA* expression, we would then be able to pinpoint which step of the nutritional response is defective in aged flies. Interestingly, it has been shown that *foxo* remains inside the nucleus even in aged fly tissues, but its target genes are different from younger fly tissues^44^. Met-dependent changes in chromatin status can be another mechanism, as SAM-dependent modification of histones and nucleic acids should be altered during early MetR^71–73^. Furthermore, it would be intriguing to explore whether the mechanism of this age-dependent transition of the nutritional response is conserved in humans.

## Acknowledgements

We would like to acknowledge Kyoto Stock Center, National Institute of Genetics, Vienna Drosophila Resource Center, and Bloomington Drosophila Stock Center for reagents. We thank Takao Fujisawa, Yoriko Akuzawa, Ayano Oi and all the members of Miur’’s laboratory and Obat’’s laboratory for technical assistance and critical advice. We thank Sa Kan Yoo and Scott Pletcher for critical reading of the manuscript. This work was supported by AMED-PRIME to F.O. under Grant Number 20gm6310011h0001 and by AMED-Project for Elucidating and Controlling Mechanisms of Aging and Longevity to M.M under Grant Number JP21gm5010001. This work was also supported by grants from the Japan Society for the Promotion of Science to F.O. under Grant Number 19H03367 and 20H05726, and to M.M. under Grant Numbers 16H06385, 21H04774 and 21K19206.

## Author Contributions

M.M. and F.O. conceived the project. H.K., J.K., H.A., R. O., T. O. and F.O. performed the experiments and analysed the data. J.J. and M. P. performed statistical analysis of lifespan curves. H.K., M.M. and F.O. wrote the initial manuscript. M.M. supervised the study. All authors approved the final manuscript.

## Materials & Correspondence

All the materials generated in this study are available upon request to F.O. or M.M.

## Methods

### *Drosophila* stocks and husbandry

Flies were reared on a standard diet containing 4% cornmeal, 4-6% yeast, 6% glucose, and 0.8% agar with 0.3% propionic acid and 0.15% nipagin. Canton-S, *w^iso^*^31^ (from Dr Alex Gould), and *w^Dah^* (from Dr Linda Partridge) flies were used as control strains. Adult flies were maintained under conditions of 25 °C and 60-65% humidity with 12 h/12 h light/dark cycles. To allow for synchronized development with constant density, embryos were collected on agar plates (2.3% agar, 1% sucrose, and 0.35% acetic acid) with live yeast paste and directly added to the standard diet in a bottle with a fixed volume. Normally, 150–200 adult flies/bottle were obtained.

The fly lines used in this study were *da-GeneSwitch* (from Dr Monnier Veronique), *NP1-Gal4, tub-Gal80^ts^*^74^, *UAS-foxo-RNAi* (BDSC, 27656)*, bmm^WT^*^43,54^*, bmm^1^*^43,54^*, ovo^D1^* (BDSC, 1309)*, UAS-lacZ* (Kyoto, 106500), *UAS-foxo* (Bloomington, 9575), and *MsrA^EY^*(Bloomington, 16671). Unless otherwise stated, female flies were used for all experiments.

### Fecundity analysis

After flies were raised on the standard yeast diet for two days after eclosion, 15 female and 15 male flies were collectively transferred to the control holidic medium or methionine-restricted medium for mating. Flies were transferred to fresh vials every three days. One week after dietary manipulation, anaesthetised flies were separated into groups of three males with three females. After 24 hours, the egg number in each vial was counted.

### Climbing ability

A negative geotaxis assay was used to quantify climbing ability. For each condition, 10-30 flies were transferred to each cuboid vial (w: 2.6 cm d: 1 cm h: 16.4 cm) using a funnel. These vials were placed in front of a light box (HOZAN, 8015-011902) inside a dark tent to shed light from behind. Flies inside were dropped to the bottom of the vial by three consecutive taps. One minute after this practice tap, the actual assay was performed and recorded using a web camera (logicool, 860-000336). The height of the flies ten seconds after the tap was measured manually. Four vials were prepared for each group.

### Dietary manipulations

The exome-matched version of a chemically defined diet, or holidic medium, was used for dietary manipulations^26,75^. Methionine restriction (MetR) is achieved by decreasing the Met concentration. To minimise the batch effect, the medium was prepared without Met and then split into two bottles, followed by the addition of Met stock solution (or just Milli-Q water for the Met(-) diet) to each bottle to make the control and MetR diets. We originally used 1 mM vs. 0.15 mM Met for MetR; however, in the middle of the study, we empirically noticed that 1 mM Met was not adequate for stably producing““well-fe”” physiological conditions. We increased the Met concentration to 1.6 mM, which was 40% of the original concentration of the holidic medium. We confirmed that all results were reproduced well by both MetR conditions. For lifespan analysis in Fig. 2j, we used original (100%) AA concentration which contains 4 mM of Met. All the ingredients used and the procedure to make the holidic medium are described in the Supplementary Data 4. The food calorie is calculated based on the content of sucrose (3.87 kcal/g), amino acids (4 kcal/g), and agar (0.031 kcal/g).

### Lifespan measurement

Adult flies that eclosed within one day were collected and maintained for an additional two days for maturation and mating on the standard diet. Flies were then sorted by sex and maintained at a fixed density (15 flies/small vial or 30 flies/large vial). For lifespan analysis, the number of dead flies was counted every three to four days when flies were transferred to fresh vials. For each lifespan curve, at least six vials were used in parallel to minimise inter-vial variation. PPG (Sigma, P7888-1G) was added to the diet to inhibit the transsulfuration pathway.

### Starvation/H_2_O_2_ resistance analysis

Female flies were transferred to a 1% agar diet (for starvation) or a 3% H_2_O_2_, 5% sucrose, 1% agar diet (for H_2_O_2_ resistance), and the number of dead flies was counted 2-3 times per day until all flies were dead.

### RNA sequencing analysis and quantitative RT‒PCR

For RNAseq analysis the guts of young or old flies upon 24 hr Met restriction, total RNA was purified from eight guts. Tissues were homogenised in Qiazol (Qiagen, 79306) using a pellet pestle (bms, BC-PES50S). RNA purification was performed using an RNeasy micro kit (Qiagen, 74004). Triplicate samples were prepared for each experimental group. The RNA samples were sent to Macrogen, where RNA sequencing was performed using an Illumina NovaSeq 6000. The paired-end sequence data were analysed as follows: a quality check of the raw reads was performed by FastQC (v0.11.9). The raw reads were filtered to remove the adaptors and low-quality bases using Trimmomatic (v0.39). Filtered reads were aligned to the *Drosophila* genome (BDGP6.22) using Hisat2 (v2.1.0). The TPM values were calculated using subread (v1.6.5). Differentially expressed genes were identified using DESeq2 (v1.26.0). RNA-sequencing data have been deposited at DDBJ with accession number DRA013585.

For 3’ RNA-seq analysis of guts after 6 hours, 2 weeks, or 4 weeks of Met restriction, 10 guts of female *w^Dah^* flies were dissected. For RNAseq analysis of abdomens upon 4 weeks of Met restriction, 10 abdomens of female *w^Dah^* flies were dissected. Triplicate samples were prepared for each experimental group. Met restriction was started on Day 5 after 3 days of feeding with the control holidic medium. Total RNA was purified using the Promega ReliaPrep RNA Tissue Miniprep kit (z6112). RNA was sent to Kazusa Genome Technologies to perform 3’ RNA-seq analysis. The cDNA library was prepared using the QuantSeq 3’ mRNA-Seq Library Prep Kit for Illumina (FWD) (LEXOGEN, 015.384). Sequencing was performed on an Illumina NextSeq 500 using the NextSeq 500/550 High Output Kit v2.5 (75 cycles) (Illumina, 20024906). Raw reads were analysed by the BlueBee Platform (LEXOGEN), which performed trimming, alignment to the *Drosophila* genome, and counting of the reads. The count data were statistically analysed by the Wald test using DESeq2. The results have been deposited in DDBJ under accession number DRA013644. For the prediction of *cis*-regulatory elements, i-*cis*Target was used^42^.

For quantitative RT‒PCR analysis, total RNA was purified from 4 tissues of female *w^Dah^* flies using the Promega ReliaPrep RNA Tissue Miniprep kit (z6112). cDNA was made from 100-400 ng of DNase-treated total RNA using a Takara PrimeScript RT Reagent Kit with gDNA Eraser (Takara bio RR047B). Quantitative PCR was performed using TB Green™ Premix Ex Taq™ (Tli RNaseH Plus) (Takara bio RR820W) and a QuantStudio 6 Flex Real Time PCR system (Thermo Fisher) using *RNA pol2* as an internal control. Primer sequences were: MsrA-Forward, gccggttcacgatgtgaatg, MsrA-Reverse, gtagcccacggtggttctc, MsrA^EY^ mutation check-Forward, agctcaaggatctgagcacc, MsrA^EY^ mutation check-Reverse, ccgtggctttggtgacattc, RNA pol2-Forward, ccttcaggagtacggctatcatct, and RNA pol2-Reverse, ccaggaagacctgagcattaatct

### Measurement of metabolites

Metabolites were quantified by ultra-performance liquid chromatography–tandem mass spectrometry (LCMS-8060, Shimadzu) based on the Primary Metabolites package ver.2 (Shimadzu)^56,57^. Four whole bodies of female flies were homogenised in 160 μL of 80% methanol containing 10 μM internal standards (methionine sulfone and 2-morpholinoethanesulfonic acid). After centrifugation at 4 °C and 20,000 × g for 5 minutes, 150 μL of the supernatant was deproteinized by mixing it with 75 μL of acetonitrile. The supernatant was placed into a pre-washed 10 kDa centrifugal device (Pall, OD010C35), and the flow-through after centrifugation at 4 °C and 14,000 × g for 5 minutes was evaporated completely using a centrifugal concentrator (TOMY, CC-105). Hemolymph was extracted according to the following protocol: A glass capillary was used to prick the side of the female abdomen, after which 10 flies were collected in a 0.5 mL tube that had been pierced at the bottom by a syringe. The resulting sample was then placed on top of a 1.5 mL tube and centrifuged at 4 °C and 8,000 × g for 5 minutes. Next, the collected hemolymph sample in the 1.5 mL tube was mixed and vortexed with 50% methanol containing 10 μM internal standards. Chloroform (250 μL) was added to the sample and vortexed. After centrifugation at 4 °C and 2,300 × g for 5 minutes, 200 μL of the supernatant was deproteinized by mixing it with 100 μL of acetonitrile. The supernatant was placed into a pre-washed 10 kDa centrifugal device (Pall, OD010C35), and the flow-through after centrifugation at 4 °C and 14,000 × g for 5 minutes was evaporated completely using a centrifugal concentrator (TOMY, CC-105). The dried and concentrated samples were resolubilised in ultrapure water and subjected to LC‒MS/MS with a PFPP column (Discovery HS F5 (2.1 mm × 150 mm, 3 μm), Sigma‒Aldrich) in a column oven at 40 °C. A gradient from solvent A (0.1% formic acid, water) to solvent B (0.1% formic acid, acetonitrile) was performed for 20 minutes. MRM method parameters were optimised by the injection and analysis of pure standards through peak integration and parameter optimisation with the use of software (Labsolutions, Shimadzu). The concentrations of metabolites were normalised by methionine sulfone and the body weight of flies or protein amount in the hemolymph. For body weight measurement, flies were anaesthetised by CO_2_ and placed on a microbalance (METTLER TOLEDO, XPR2).

### Lipid staining

Oil red O staining was performed as described below. Oil red O (0.5%, Fujifilm Wako, 154-02072) in 100% isopropanol was made as a stock solution and mixed by vortexing. Guts were dissected and fixed with 4% paraformaldehyde (Nacalai Tesque, 09154-85) for 20 min. The guts were washed with 1× PBS twice. The oil red O stock solution was freshly mixed with Milli-Q water at a ratio of 3:2 by vortexing and filtrated using a 0.45 µm filter. The gut samples were incubated with 500 μL of fresh oil red O solution and incubated for 30 min. After washing with Milli-Q water twice, the guts were mounted in 80% glycerol. Images were obtained by a Leica MZ10F and a Zeiss Axio Virt.A1.

LipidTOX staining was performed as described below. Guts were dissected and fixed with 4% paraformaldehyde (Nacalai Tesque, 09154-85) for 20 min. The guts were washed with PBST (0.1% Triton X-100) three times and incubated with LipidTOX Deep Red neutral lipid stain (Invitrogen, H34477, 1:250) for two hours. After washing with PBST (0.1% Triton X-100), the guts were mounted in 80% glycerol. Images were obtained by a Zeiss Axio Virt.A1.

### Single-cell transcriptome analysis

Guts of anaesthetised adult female flies were dissected in 1 × PBS. The proventriculus and hindgut were carefully removed. The dissected midguts were placed in ice-cold Schneider’s medium (without FBS or penicillin/streptomycin). Collected midguts (100 midguts/sample) were washed briefly with PBS and transferred to a 1.5 mL Eppendorf tube containing 5 mg/mL elastase/PBS solution (SIGMA, E0258). The midgut solution was incubated on a shaker at 27 °C and 1,000 rpm for 25 min with pipetting (40 times) every five minutes. Pipette tips were precoated with elastase solution to prevent the midgut from sticking to them. The dissociation reaction was stopped by adding 400 μL of Schneider’s medium. The cell suspension was passed through a 100 μm cell strainer. The flowthrough was centrifuged at 600 × g for 20 minutes, and the supernatant was discarded. The cell pellet was resuspended in 100 μL of Schneider’s medium. Cell number and viability were assessed by trypan blue staining using a haemocytometer. The samples were then sent to Genble, Inc. Library preparation, sequencing, and data processing were performed as follows.

A single-cell suspension was obtained by filtering the cell suspension with a 40 μm cell strainer. Isolated single cells were loaded onto the 10x Chromium Controller (10x Genomics). Single-cell cDNA synthesis, amplification and sequencing library creation were performed by using the Single Cell 3’ Library kit v3.1 following the manufacturer’s protocol. The libraries were sequenced on a NovaSeq 6000 (Illumina) platform with the following sequencing parameters: 28 bp read 1, 8 bp index 1, 91 bp read 2. Sequenced reads were subjected to demultiplexing, alignment, barcode counting, UMI counting, and filtering using Cell Ranger v.5.0.1. The *Drosophila melanogaster* genome (BDGP6.28) was used as a reference to align reads. Subsequent analysis was performed in R (version 3.6.0) using the package Seurat (version 3.1.0). Genes expressed in fewer than three cells were removed. The cells that expressed less than 200 genes or that expressed more than 3,000 genes were removed. We also removed the cells that had more than 25% of mitochondrial-associated genes among their expressed genes and more than 15,000 UMI counts. The UMI counts were log normalised (scale factor = 10,000). We regressed out the S scores, G2/M scores, and the percentage of mitochondrial-associated genes during data scaling. Dimensionality reduction was performed using principal component analysis (PCA). The first 50 principal components were used to cluster the cells based on the shared nearest neighbours (SNN) algorithm and visualised in two dimensions by uniform manifold approximation and projection (UMAP). Differential gene expression analysis was performed using MAST^76^ in the Seurat package with min.pct at 0.2. The dataset was deposited in the Gene Expression Omnibus database under accession number GSE198149.

### Heatmap, PCA, and dot plot analyses for transcriptome datasets

For the heatmap analysis and the PCA, DEGs expressed in more than two clusters in cells classified as ISC or EB were assessed in the progenitor cell cluster. DEGs expressed in more than six clusters in cells classified as aEC, mEC, copper cell/iron cell, pEC, or LFC were assessed in the enterocyte cluster. DEGs expressed in more than four clusters in cells classified as EE were assessed in the enteroendocrine cell cluster. PCA was performed using the mixOmics package (3.14)^58^. A dot plot was generated using the ggplot2 package (3.3.5).

### Statistical analysis

Statistical analysis was performed using GraphPad Prism 8/9. The sample numbers were determined empirically. All data points were biological, not technical, replicates. A two-tailed Student’s *t* test was used to test between samples. One-way ANOVA with Holm-Šídák’s multiple comparison test was used to test among groups. One-way ANOVA with Dunnett’s multiple comparison test was used to test samples against controls. All experimental results were tested at least twice to confirm their reproducibility. Bar graphs are drawn as the mean and SEM. OASIS 2 was used to perform log-rank test for lifespan analysis^77^. Cox PH analysis was performed using R^78^. The Cox Proportional-Hazards Model (‘coxph’) function of the “Survival” package^79^ was used to fit the model and the ‘Anova’ function of the “car” package^80^ to assess lifespan differences between diets and timings of dietary treatment or between diets and genotype, as well as diet by timing interactions or diet by genotype interactions. The code is in a Supplementary Data 5.

### Data availability

All relevant data are available from the authors upon request. The NGS data are available under accession numbers DRA013585, DRA013644, and GSE198149.

### Code availability

No custom codes were used during this study.

---

**Supplementary Fig. 1.**
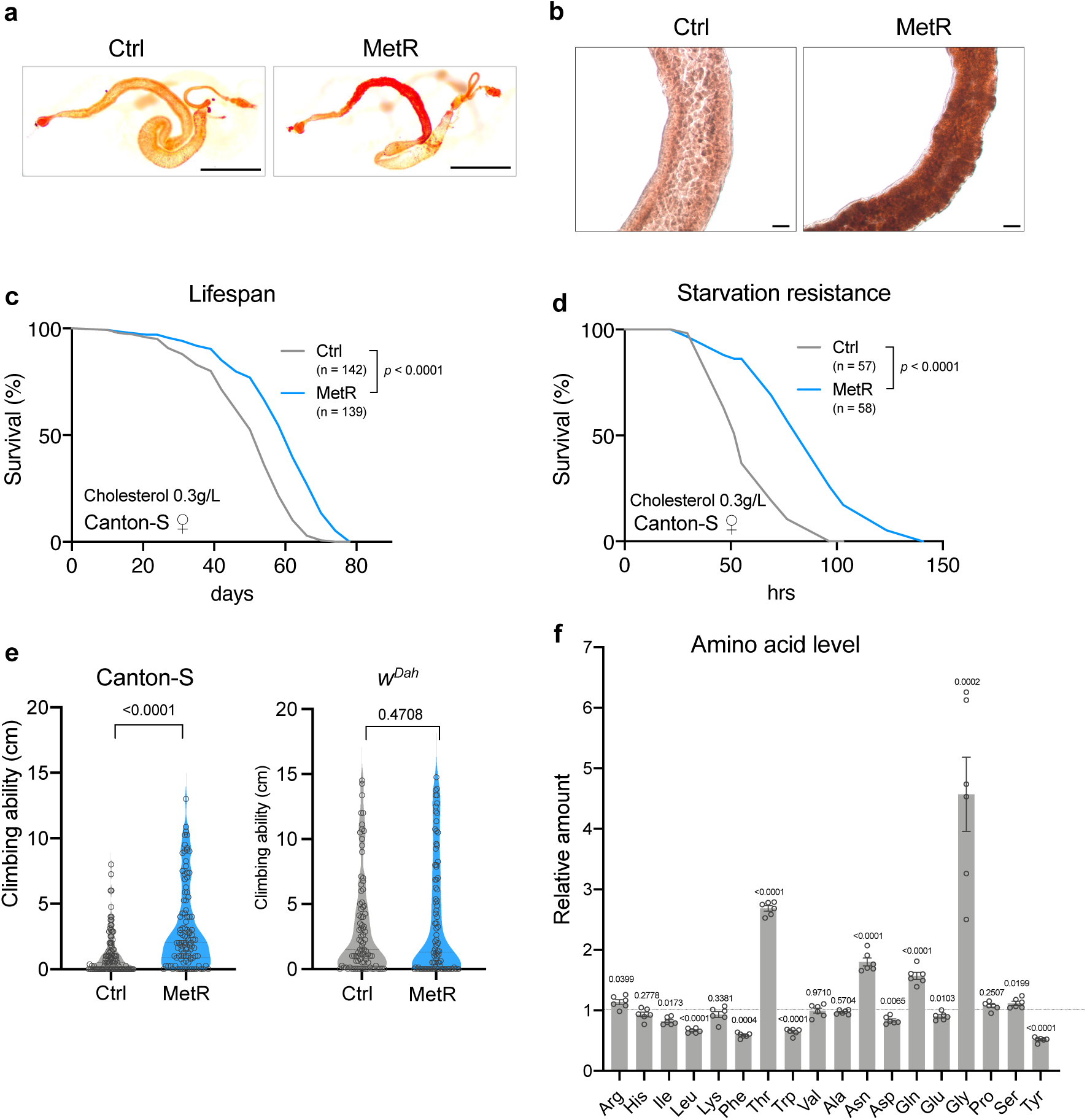
Phenotypic analysis of methionine-restricted flies. **a, b,** Oil red O staining of female guts of Canton-S flies fed with or without a methionine-restricted diet for nine days. Whole gut (**a**) or magnified view of the anterior midgut (**b**). Scale bar: 1 mm (**a**), or 100 μm (**b**). **c**, Lifespans of female Canton-S flies fed with or without a methionine-restricted diet that contained three times as much cholesterol. Sample sizes (n) are shown in the figure. For the statistics, a log-rank test was used. **d,** Survivability of female Canton-S flies upon complete starvation after feeding with or without a methionine-restricted diet that contained three times as much cholesterol for a week. Sample sizes (n) are shown in the figure. For the statistics, a log-rank test was used. **e,** Climbing abilities of female Canton-S or *w^Dah^* flies fed with or without a methionine-restricted diet for four weeks. **f,** Quantification of amino acids other than methionine in female *w^Dah^* flies upon methionine restriction. The relative amount of each amino acid upon methionine restriction compared to the control diet is shown. n = 6. For the statistics, a two-tailed Student’s *t* test was used. For the graph, the mean and SEM are shown. Data points indicate biological replicates.

**Supplementary Fig. 2.**
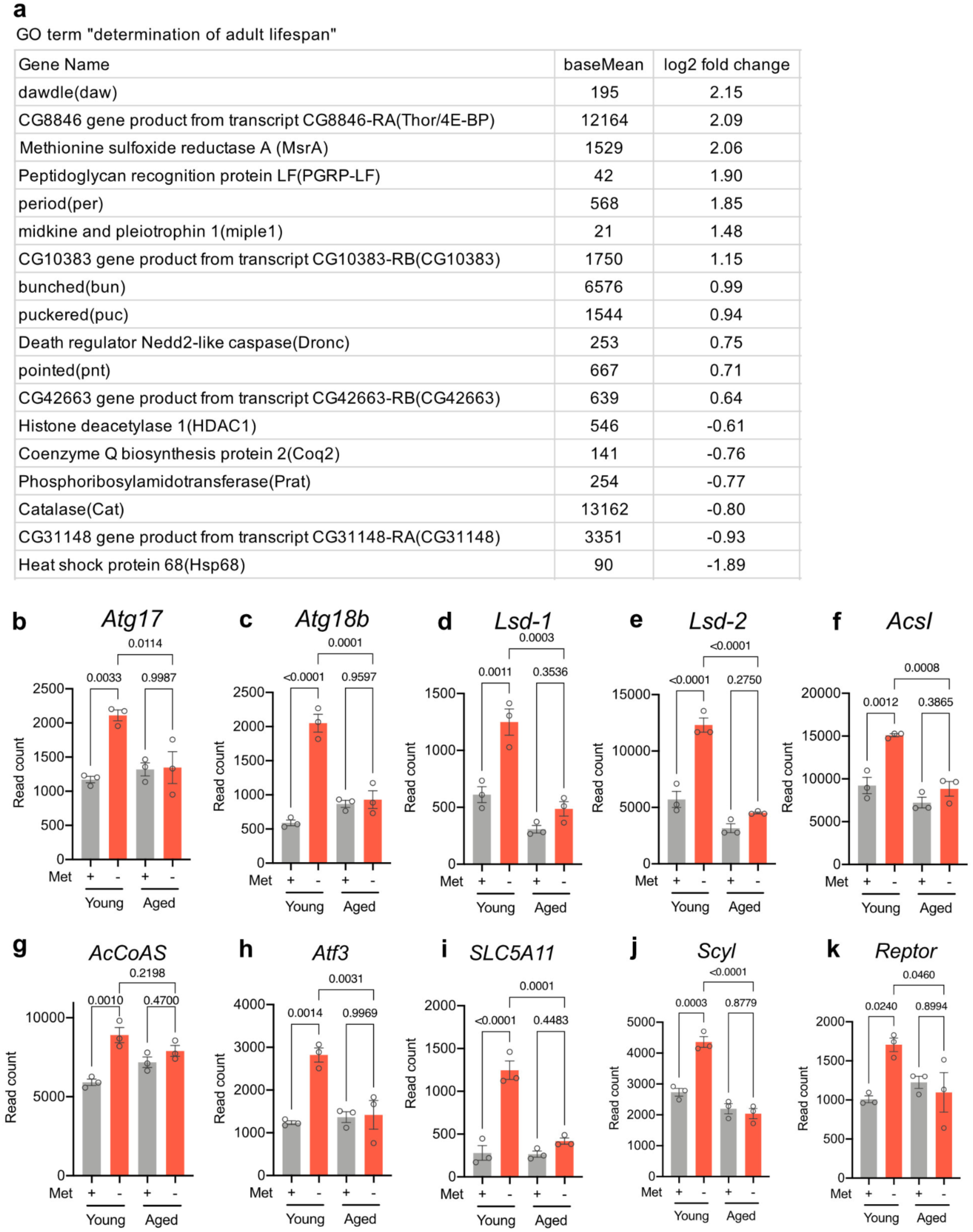
Transcriptomic responses to methionine restriction in young and aged flies. **a,** List of genes termed “determination of adult lifespan” by Gene Ontology analysis. **B-k,** Read counts of genes induced by methionine restriction at a young age from the RNAseq analysis. N = 3. For the statistics, one-way ANOVA with Holm-Šídák’s multiple comparison test was used. For all graphs, the mean and SEM are shown. Data points indicate biological replicates.

**Supplementary Fig. 3.**
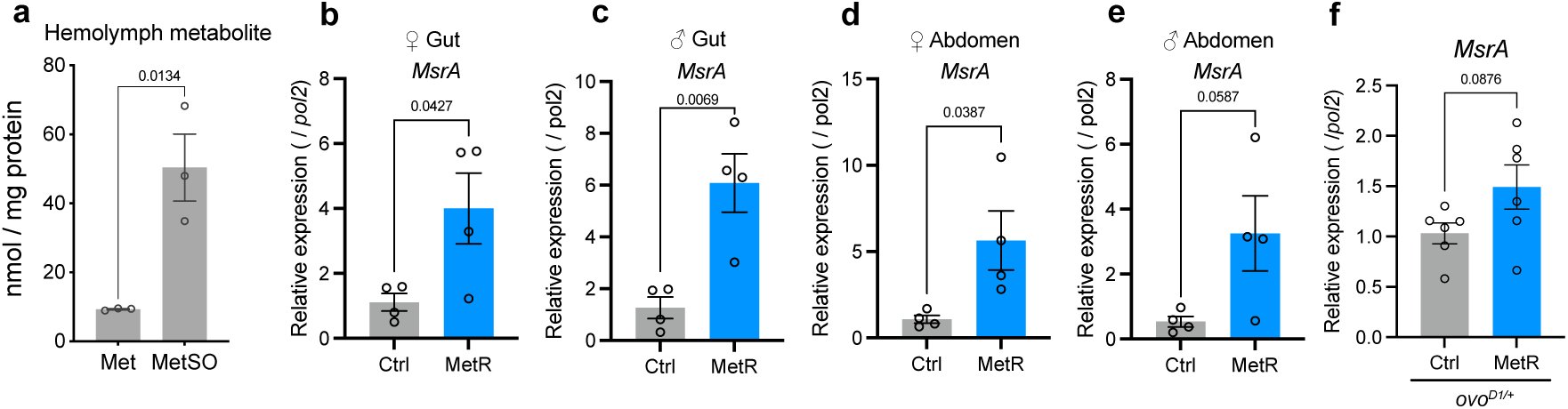
*MsrA* induction upon methionine restriction. **a,** Quantification of Met and MetSO in the hemolymph of female Canton-S flies that were fed with a standard yeast-based diet for four days post-eclosion. n = 3. For the statistics, a two-tailed Student’s *t* test was used. **b-e,** Quantitative RT‒PCR analysis of *MsrA* expression levels in female guts (**b**), male guts (**c**), female abdomens (**d**) and male abdomens (**e**) of Canton-S flies fed with or without a methionine-restricted diet for three days. n = 4. For the statistics, a two-tailed Student’s *t* test was used. **f**, Quantitative RT‒ PCR analysis of *MsrA* expression in female guts of *ovo^D1/+^* fed with or without a methionine-restricted diet for three days. n = 6. For the statistics, a two-tailed Student’s *t* test was used. For all graphs, the mean and SEM are shown. Data points indicate biological replicates.

**Supplementary Fig. 4.**
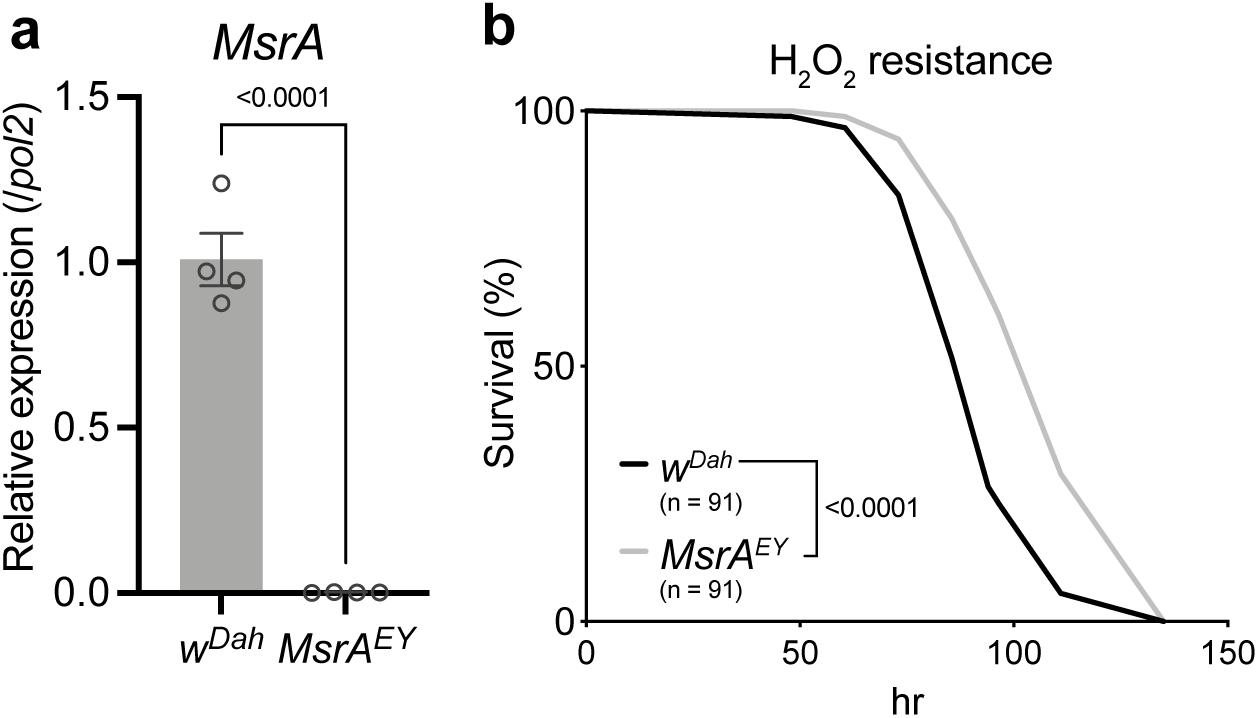
Characterisation of the *MsrA* mutant. **a,** Quantitative RT‒PCR analysis of *MsrA* in the whole bodies of *w^Dah^* or *MsrA^EY^* flies. n = 4. For the statistics, a two-tailed Student’s *t* test was used. **B**, Survivability of female flies of *w^Dah^* or *MsrA^EY^* upon 3% H_2_O_2_ treatment. Sample sizes (n) are shown in the figure. For the statistics, a log-rank test was used. For the graph, the mean and SEM are shown. Data points indicate biological replicates.

**Supplementary Fig. 5.**
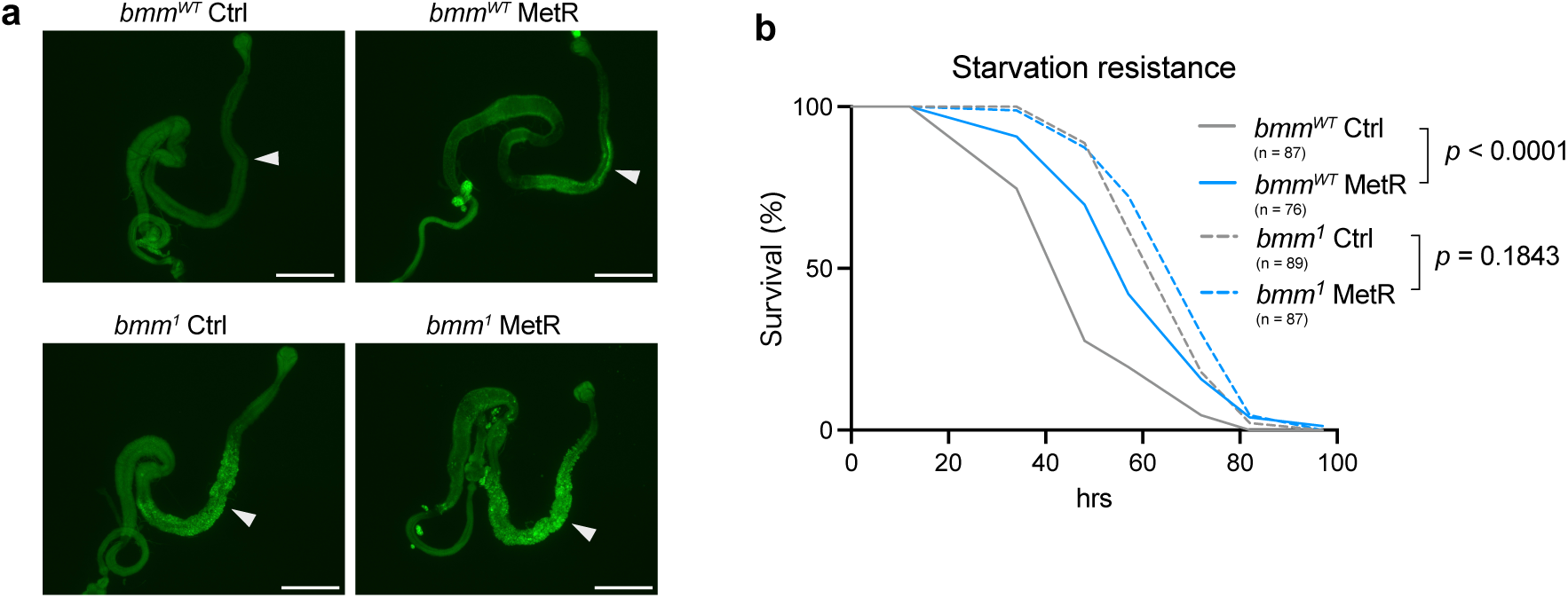
Contribution of lipid metabolism to lifespan extension upon methionine restriction. **a,** Lipid staining of the female guts of *bmm^wt^* and *bmm^1^* flies fed with or without a methionine-restricted diet for one week using LipixTOX. Scale bar: 1 mm. Arrowheads indicate lipid accumulation. **b,** Survivability of female *bmm^wt^* and *bmm^1^*flies upon complete starvation after feeding with or without a methionine-restricted diet for one week. Sample sizes (n) are shown in the figure. For the statistics, a log-rank test was used.

**Supplementary Fig. 6.**
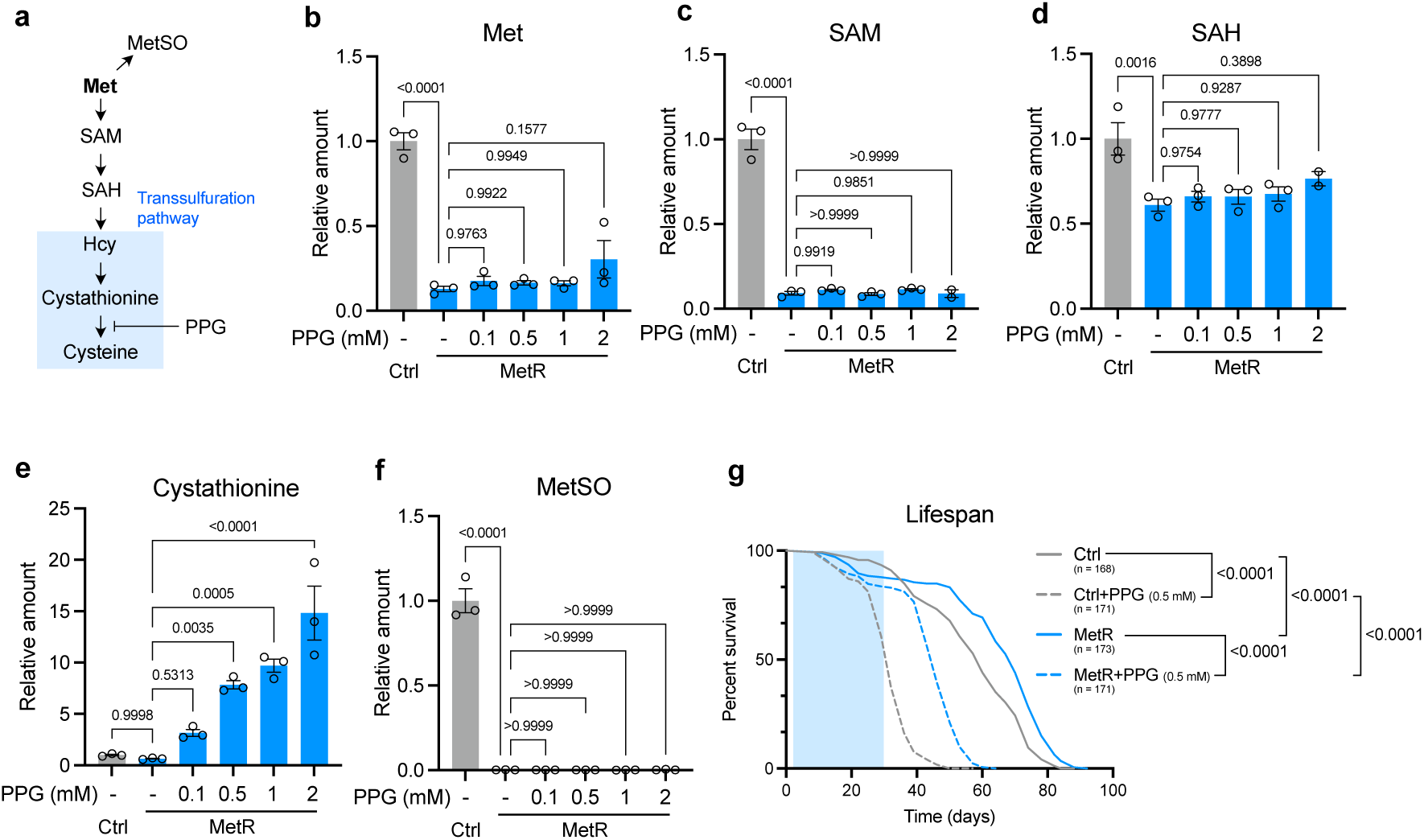
Inhibition of Transsulfuration pathway did not abrogate lifespan extension upon methionine restriction. **a,** Methionine metabolic and transsulfuration pathways, which can be inhibited by propargylglycine (PPG). **b-f,** Quantification of methionine metabolites and their oxidative products upon methionine restriction. n = 3. For the statistics, one-way ANOVA with Holm-Šídák’s multiple comparison test was used. **g,** Lifespans of female Canton-S flies fed with or without a methionine-restricted diet supplemented with 0.5 mM PPG. Sample sizes (n) are shown in the figure. For the statistics, a log-rank test was used. For all graphs, the mean and SEM are shown. Data points indicate biological replicates.

**Supplementary Fig. 7.**
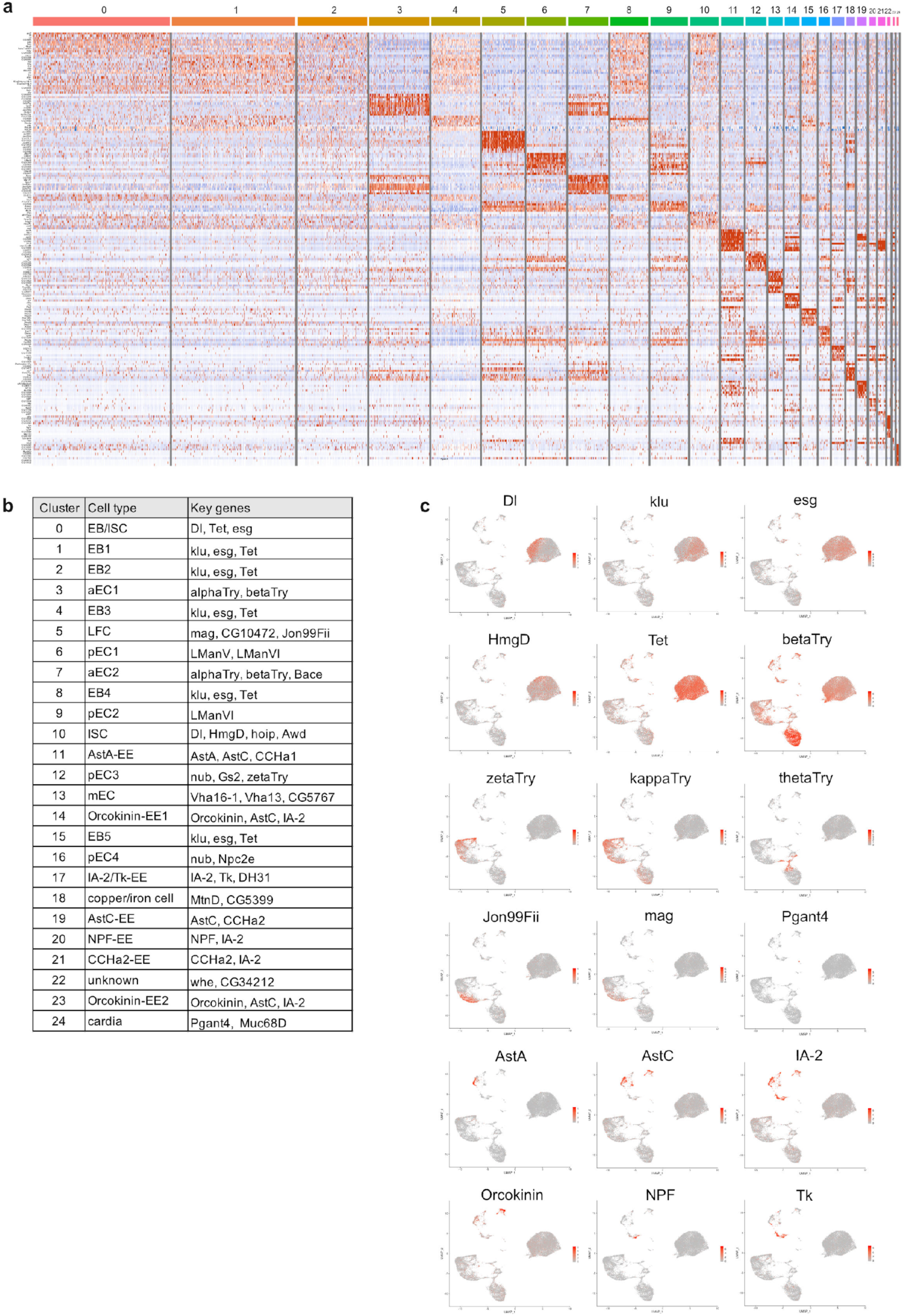
Cell type clustering analysis and marker gene expression in each cluster. **a,** Heatmap of clustered marker genes from the results of single-cell RNAseq of the *Drosophila* female midgut. **b,** Cell type names in clusters. **c,** UMAP plot of characteristic marker genes.

**Supplementary Fig. 8.**
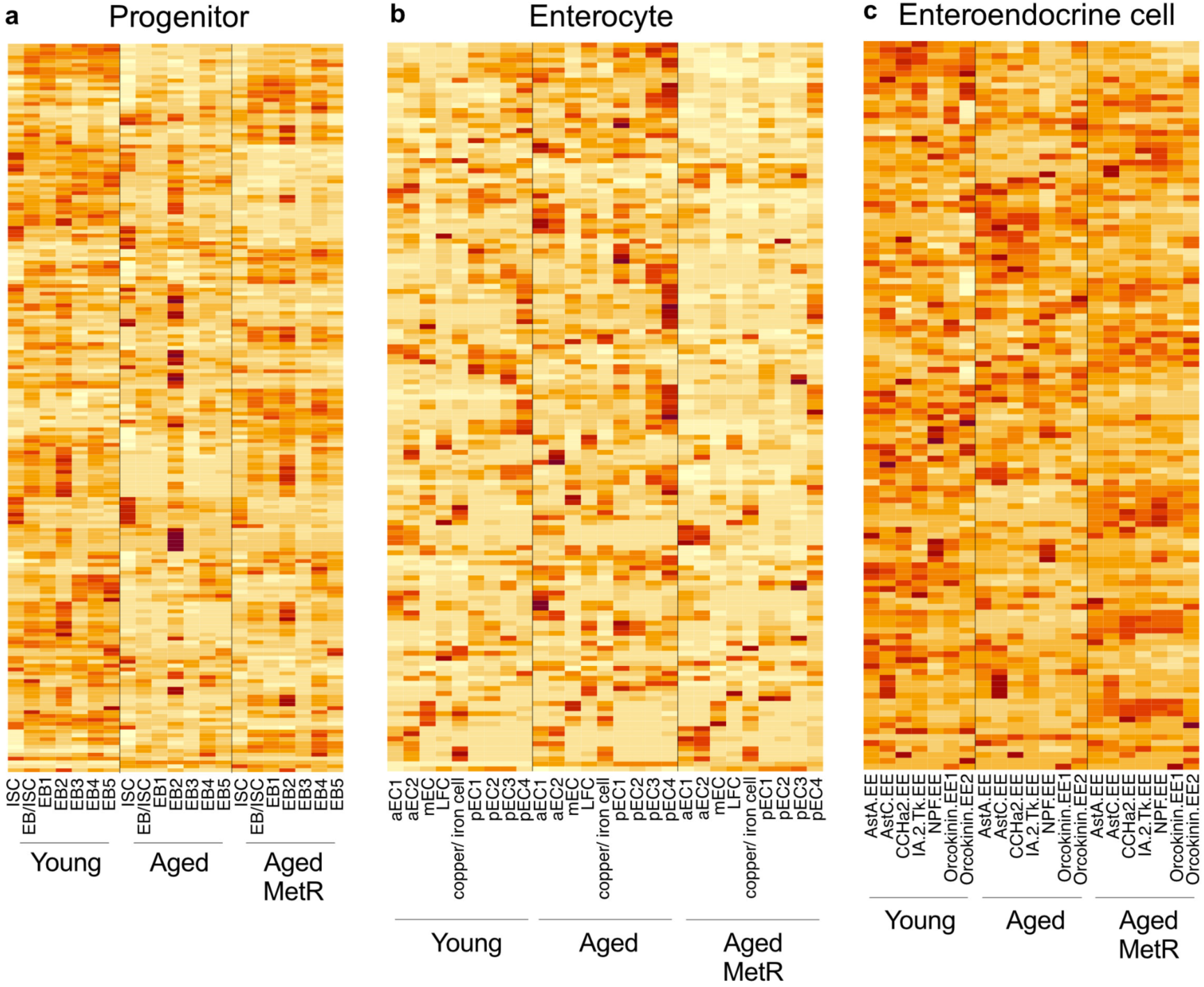
Heatmap analysis of DEGs from single-cell RNAseq. **a-c,** Heatmap of DEGs in each cell type classified as progenitor cells (**a**), enterocytes (**b**), and enteroendocrine cells (**c**) upon methionine restriction with ageing.

**Supplementary Fig. 9.**
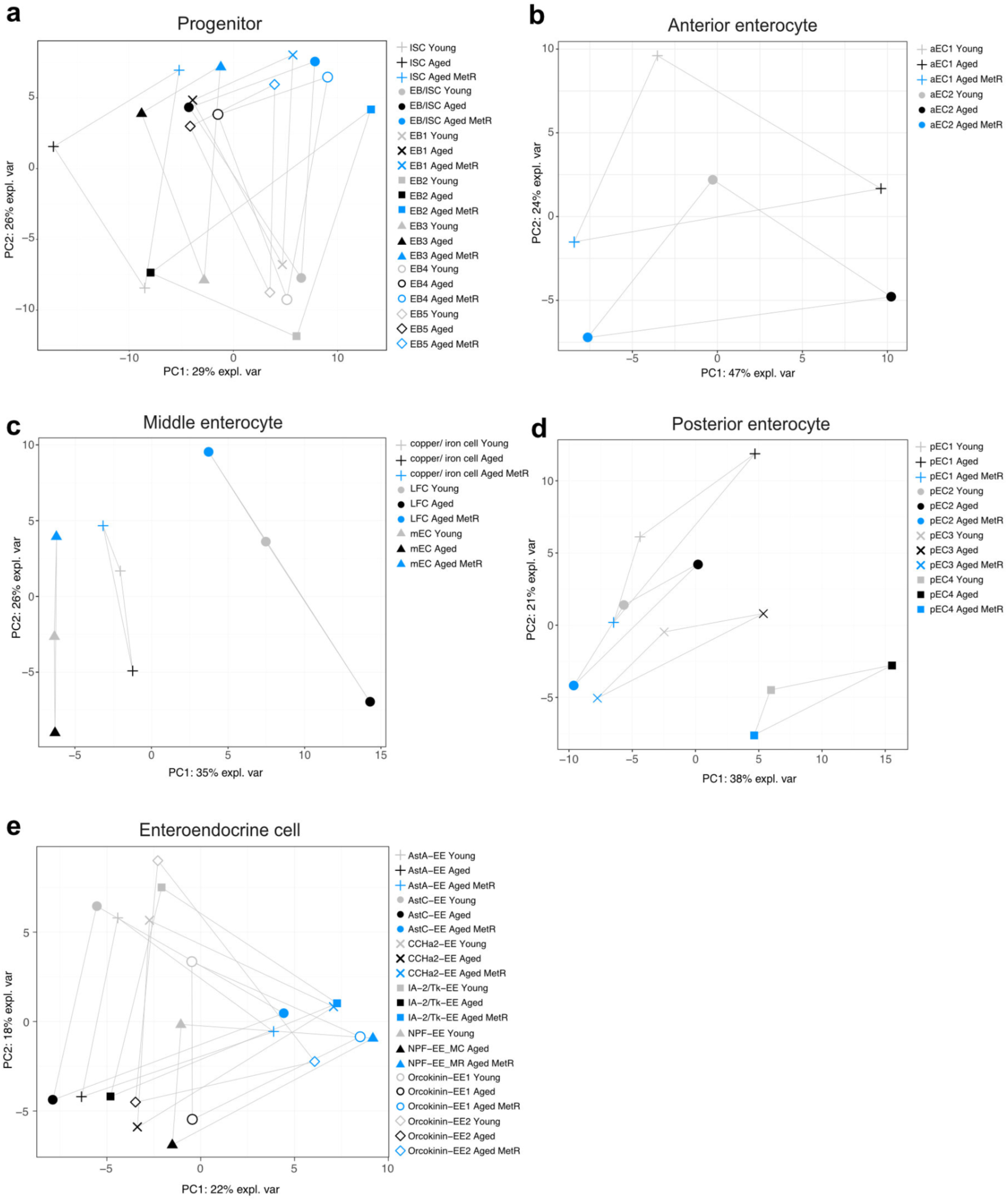
PCA of single-cell RNAseq results upon methionine restriction. **a-e,** PCA of DEGs in each cell type classified as progenitor cells (**a**), anterior enterocytes (**b**), middle enterocytes (**c**), posterior enterocytes (**d**), and enteroendocrine cells (**c**) upon methionine restriction with ageing.

**Supplementary Table 1.** Cox PH analysis for *w^iso31^* and *w^Dah^* female flies.

| *Dependent variable:* |  |  |
| --- | --- | --- |
|  | Age (days) |  |
| | $w^{iso31}$ | $w^{Dah}$ |
|  | coefficient (SE) | coefficient (SE) |
| Timing of diet treatment | -0.156 (0.125) | -0.048 (0.119) |
| MetR diet | -0.753<sup>***</sup> (0.130) | -0.524<sup>***</sup> (0.120) |
| Timing of diet treatment: MetR diet | 0.589<sup>***</sup> (0.178) | 0.191 (0.167) |
| Observations | 528 | 576 |
| Score (Logrank) Test (df = 3) | 38.854<sup>***</sup> | 27.841<sup>***</sup> |
| *** $p < 0.001$ | | |

**Supplementary Table 2.** Cox PH analysis for *MsrA^EY^* flies compared to *w^Dah^* flies.

|  | *Dependent variable:* |
| --- | --- |
|  | Age (days) |
|  | coefficient (SE) |
| Genotype | -0.367*** (0.109) |
| MetR diet | -1.240*** (0.110) |
| Genotype: MetR diet | 1.264*** (0.153) |
| Observations | 703 |
| Score (Logrank) Test (df = 3) | 143.7*** |
| | *** $p < 0.001$ |

**Supplementary Table 3.** Cox PH analysis for *bmm^1^* flies compared to *bmm^WT^* flies.

| *Dependent variable:* |  |
| --- | --- |
|  | Age (days) |
|  | coefficient (SE) |
| Genotype | 0.946*** (0.117) |
| MetR diet | -0.790*** (0.116) |
| Genotype: MetR diet | -0.695*** (0.160) |
| Observations | 663 |
| Score (Logrank) Test (df = 3) | 248*** |
| *** $p < 0.001$ | |

